# Physical association of functionally antagonistic enzymes: KDM5A interacts with MLLs to regulate gene expression in a promoter specific manner facilitating EMT and pluripotency

**DOI:** 10.1101/2021.12.03.471073

**Authors:** R Kirtana, Soumen Manna, Samir Kumar Patra

## Abstract

Differential expression of genes involved in physiological processes are a collaborative outcome of interactions among signalling molecules, downstream effectors and epigenetic modifiers, which together dictate the regulation of genes in response to specific stimuli. MLLs and KDM5A are functionally antagonistic proteins as one acts as writer and the other as eraser of the active chromatin mark, i.e., H3K4me3. KDM5A promotes EMT by occupying promoters of both epithelial and mesenchymal markers. Through this work, it is illustrated that when bound to E-cadherin promoter, KDM5A acts as a classical repressor by demethylating H3K4me3, but on mesenchymal marker promoters, it acts as a transcriptional activator by inhibiting the activity of HDACs and increasing H3K18ac. Further it is demonstrated that KDM5A occupancy enhances either MLL1 or MLL2 by physically interacting with them and that signalling pathways regulate the enzymatic activity of KDM5A probably by phosphorylation. When not active, KDM5A signals for MLL occupancy, a mechanism that can be called epigenetic signalling.

## Introduction

Manifestation of a physiological process is an outcome of certain stimulus that triggers signalling pathways leading to activation of transcriptional factors which further couple with epigenetic modifiers to setup a desired chromatin state at target gene TSS (1). The processes of EMT and pluripotency are governed by a fine-tuned balance of many epigenetic modifications like DNA methylation, histone acetylation and methylation and also by miRNAs (2).

H3K4me3 is generally associated with transcriptionally active chromatin (3,4) but at bivalent promoters, it corresponds to poised state due to the presence of H3K27me3, and is primed to active configuration only when the repressive mark is removed (5). This active mark is catalysed by highly conserved Set domain containing proteins i.e., SET1A/B, MLL1-5 in humans. Structurally, MLL1 and MLL2 possess similar domain configuration – with AT rich, PHD, BRD, FYR and SET domains and functionally share the core subunits (WDR5, RBBP5, ASH2L and DPY30) to form an enzymatically active complex. Mll1 homozygous deletions in mice are embryonic lethal and heterozygous embryos show retarded growth and hematopoietic defects. MLL2 is essential during embryonic development but is dispensable after E11.5 in mouse models and Mll2^-/-^ ES cells show high apoptosis and skewed differentiation along with prolonged Oct4 expression. At bivalent promoters, MLL2 is the main methyltransferase and its loss leads to increased PRC2 complex occupancy (6,7,8).

Demethylation of H3K4me3 is catalysed by two families of proteins – LSD (FAD dependent oxidases) and JARID (alpha-KG dependent hydroxylases). KDM5A^-/-^ are viable owing to the compensatory role of other family members and earlier work suggests that it regulates stemness, differentiation, cell cycle progression and mitochondrial functions by interacting with other epigenetic regulators like EZH2 and HDACs (9).

Multiple parameters trigger epithelial cells to attain a reversible fibroblast like morphology, a process broadly classified as EMT. Type-III EMT, the one associated with cancer cells, can exist in a hybrid epithelial/mesenchymal phenotype with internalized epithelial proteins, as opposed to the classical EMT which is manifested by downregulation of epithelial proteins (10,11). Of the many factors that influence this multistep process, two important signalling pathways – the ones mostly abrogated in cancer cells and which show bi-directional dependence are calcium signalling and integrin-FAK pathway. The higher calcium concentrations at the rear end aid in dissolving cell adhesions and the new focal adhesions assembling at front end activate L-type calcium channels that further stabilize filopodia and promote maturation of focal adhesions (12,13).

The epigenetic regulation of EMT is a less understood and paradoxical area with many gaps about the pivotal mechanism - where some reports show MLL1 expression to enhance metastasis by increasing Zeb1 expression or by interacting with B-catenin (14) and other studies show that LSD1 demethylates E-cadherin promoter thereby repressing its expression and promoting EMT, with contradictory reports on LSD1 supressing invasiveness and metastasis in breast cancer cells (15,16). KDM5A mediates EMT in renal, lung and ovarian cancers (9), but these studies do not provide proper clues regarding the mechanism of regulation.

Following EMT, many metastatic tumours undergo the reverse process i.e., MET which potentiates stem cell like properties to colonize the new distal niches after metastasis (17). Defined as indefinite self-renewal capability, pluripotency is the ability to initiate/develop into all three germ layers of an embryo and from a cancer perspective, pluripotency induces unlimited self-renewal ability via enhanced expression of transcription factors like OCT4, SOX2, NANOG and KLF4 which potentiate expression of CSC-related molecules like CD133, CD44 and ABCGs leading to tumorigenic/aggressive phenotypes (18). Many studies validated the overexpression of one or more of these transcription factors in a cancer tissue-dependent manner. Of these, NANOG overexpression in particular enhanced FAK levels, which in-turn interacts and phosphorylates NANOG protein in the nucleus, thereby aiding in filopodia formation and cell invasion, a phenomenon that were interfered either by FAK inhibition or by mutant NANOG proteins that were unable to bind FAK (19). Recently, the stem cell niche of ESCs was shown to be maintained by secreting extracellular vesicles that activated FAK and enhanced pluripotency (20).

Pluripotency associated genes are induced by the active mark H3K4me3 and some studies show that pluripotency inducers (O4I3 - Oct4 inducing compound 3) promote human iPSC formation from human primary fibroblasts by increasing H3K4me3 via inhibiting KDM5A but not KDM5B activity (21). Contradicting the previous report, mouse neural stem cells treated with VPA (a HDACi) showed enhanced OCT4 expression with increased KDM5A occupancy on its promoter, inferring that KDM5A contribute to OCT4 activation in NSC (22), and that KDM5A expression maintains NPC in undifferentiated state by repressing GFAP expression and inhibiting astrogenesis (23). As KDM5A functions in a tissue specific manner and this study is intended to explore how KDM5A acts as either a co-activator or co-repression on the same gene promoter and understand if KDM5A exhibits its activity via interactions with other epigenetic modifiers to be an activator or repressor.

## Materials and methods

Cells used along with their growth conditions, preliminary experimental setup and some confirmatory data are provided in supplementary information.

### Immunoblotting

Whole cell lysate prepared by RIPA lysis buffer (sigma –R0278-50ML) by following the protocol adapted from (24) and the proteins were electrophoresed on 8-12% SDS-PAGE gels depending on target proteins analysed, resolved and transferred to nitrocellulose membrane (Axiva – 160300RI), blocked with 5% skim milk for an hour. All primary antibodies were prepared in 1% BSA (Himedia-MB083) in PBST and the blots were probed with primary antibodies at 4°C overnight. Following three washes with PBST, the blots were incubated with either anti-rabbit (Invitrogen-65-6120) or anti-mouse (Santa Cruz – SC516102) secondary antibody for an hour. The blots were then washed thrice using PBST, visualized using ECL chemiluminescence detection system (Thermo-34580). Details of the all the primary antibodies used in this study are provided in supplementary information (refer to section – antibodies used in this study).

### Chromatin immunoprecipitation

ChIP was performed both manually and by using a kit (Sigma-Imprint chromatin immunoprecipitation kit – CHP1). When using kit, manufacturer’s instructions were followed and manually, the protocol was adapted from (25) and modified. Following the required treatments, cells were washed with PBS, fixed with 5mL of 1% formaldehyde for 10 minutes on a rocker. The reaction was quenched using 1.25M glycine (0.5ml) for 5 minutes at room temperature. Cells were pelleted, washed with PBS and lysed (on ice) for 10 minutes in 300uL of ChIP lysis buffer, followed by sonication (0.3 mm probe sonicator) on ice for 10 minutes on 30s ON-30s OFF cycles at 30% efficiency and diluted using equal amount of ChIP dilution buffer. 100uL of this diluted chromatin was incubated with 1-3ug of primary antibody overnight at 4°C, followed by addition of A/G agarose beads (Santacruz – SC-2003) and agitated for 4hrs at 4°C. The beads with immunoprecipitated complexes were washed 3 times with low salt buffer, once with high salt buffer, once with LiCl buffer and finally with TE buffer. Chromatin elution was performed using fresh elution for 1hr at room temperature, followed by crosslink reversal (using 5M NaCl) at 65°C for 3hrs followed by RNase and proteinase K treatments. The DNA was purified using Quigen PCR purification kit. Compositions of all the buffers, sequence of the primers used and %INPUT calculations are provided in supplementary information.

### Co-immunoprecipitation and western blotting

Whole cell lysates made in RIPA buffer were used for co-immunoprecipitation. 100-150ug cell lysate were incubated with 1-3 ug antibodies – anti-KDM5A, anti-MLL1 and anti-MLL2 (see supplementary information for catalog numbers) overnight at 4°C under agitation followed by addition of 30uL A/G beads (SC-2003) and agitated for another 4hrs at 4°C. The beads containing immunoprecipitated complexes were was thrice with RIPA buffer, and eluted in 40ul of SDS-loading buffer by heating for 5 minutes. The samples were analysed for protein interactions by western blotting.

### Bacterial protein purification for pulldown experiments

GST-tagged KDM5A deletion constructs (D1-D5; borrowed from Dr. Shweta Tyagi) (26) were expressed in BL21-DE3 and induced using 0.1-0.2 mM IPTG for 8hrs at 22°C. The cell pellet collected was lysed in lysis buffer (26) and sonicated, incubated with glutathione agarose beads (sigma-G4510) at 4°C for 4-6hrs. following protein quantification by SDS-PAGE, GST-bound protein was incubated with equal amount of whole cell lysates from U87MG and HaCaT for 6-8hrs. The bead bound complexes were washed thrice with RIPA buffer, eluted in SDS-loading buffer and electrophoresed to detect interactions.

### Statistical analysis

Either student’s ‘t’ test or two-way ANOVA followed by Bonferroni correction were employed to test the significance, and the test employed along with the p-values and their inference is mentioned in figure legends.

## Results

### KDM5A supresses stemness and E-cadherin expression in a demethylase dependent manner

EMT is a complex process with stages involving shedding of epithelial markers, loss of polarity, degradation of basement membrane, remodelling of cytoskeleton and adhesion junctions leading to acquisition of mesenchymal cell phenotype that can migrate and colonize at distinct sites (10). Several studies have earlier reported that the histone demethylase KDM5A induces EMT in ovarian, lung and renal carcinoma (27,28,29), but no detailed mechanism was elucidated on how KDM5A regulates this multistage process or sets control over regulation of genes of a particular stage. To understand how KDM5A regulates genes related to the process of EMT, depletion (using siRNA) and overexpressed (pcDNA-SFB tagged-RBP2 plasmid) of KDM5A in HeLa, HaCaT and PC3 cell lines (Supplementary figure S1), and analysed the expression of EMT markers at transcript and protein level. Our results infer that KDM5A knockdown enhanced E-cadherin expression and downregulated mesenchymal markers – Vimentin, Snail, Slug, Zeb1 and Twist1 (Figure 1A-C, Figure 2A). This observation was reversed on overexpressing KDM5A, proving that KDM5A acts as H3K4me3 demethylase on E-cadherin promoter thereby reducing its expression when overexpressed. To confirm this, Chromatin immunoprecipitation (in HaCaT cell line) was employed using the same experimental setup and it was observed that KDM5A knockdown lead to enhanced MLL1 and MLL2 occupancy on E-cadherin promoter (figure 3.B&C), associated with increase in H3K4me3 (figure 3D) which led to increase in its expression. On the other hand, overexpressing KDM5A lead to enhanced MLL2 occupancy (figure 3C) on E-cadherin promoter, but MLL1 and H3K4me3 decreased (figure 3.B&D), which explains the downregulation of E-cadherin. From these ChIP experiments, it was understood that presence of MLL1 and absence of KDM5A dictates E-cadherin expression. To support the above observations, MLL1 knockdown was performed which led to enhanced KDM5A and MLL2 occupancy but reduced H3K4me3 (figure 3), confirming our previous notion that KDM5A and MLL1 act on E-cadherin promoter by displacing each other.

**Figure 1:**
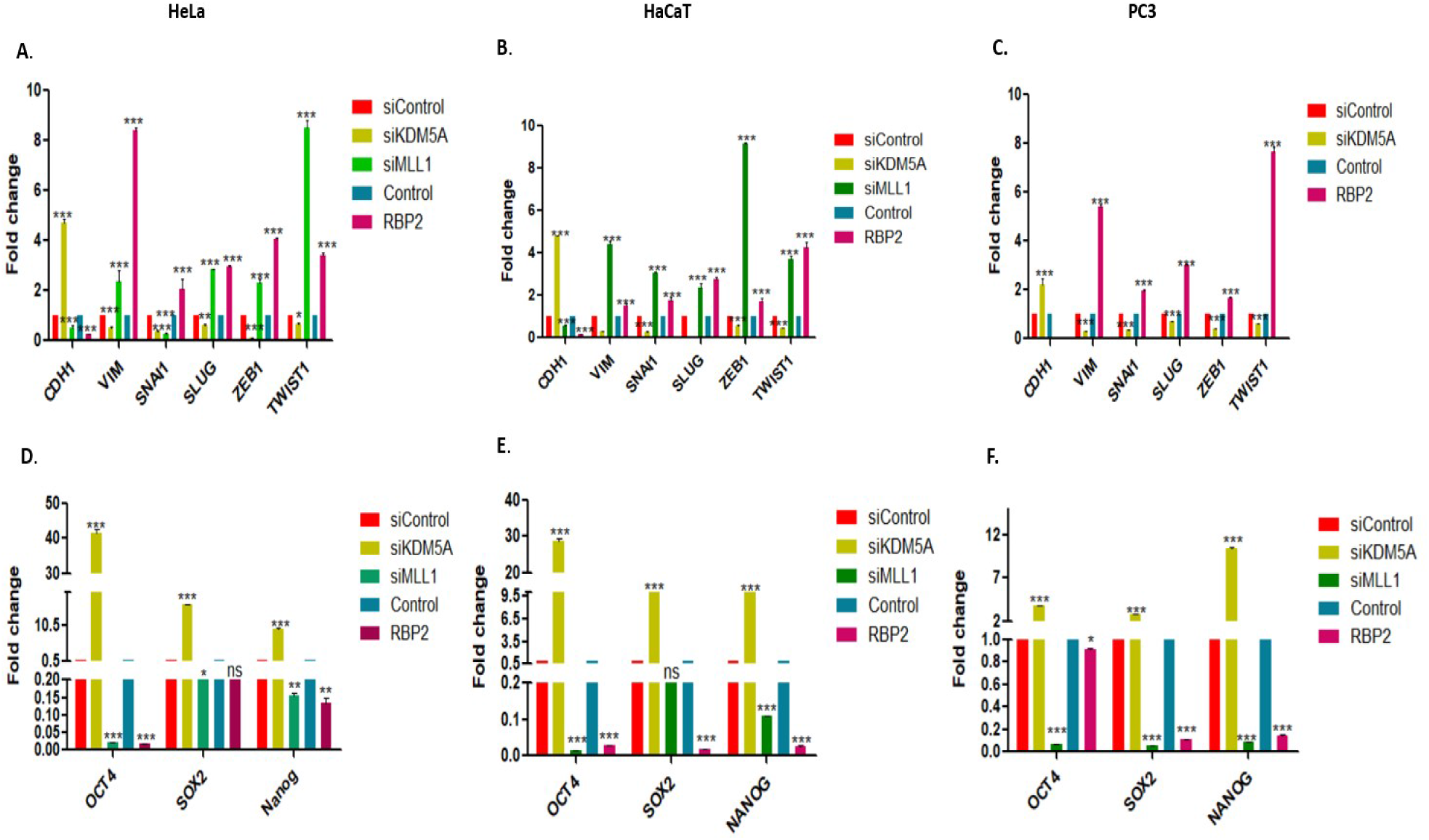
KDM5A and MLL1 regulate the expression of EMT (A, B, C) and pluripotency related genes (D, E, F). HeLa (A, D), HaCaT (B, E) and PC3 (C, F) cell lines treated with siRNA against KDM5A, orMLL1 and transfected with KDM5A overexpression construct, with either negative control siRNA or normal untreated cells used as control, followed by total RNA extraction, cDNA synthesis and quantitative PCR was performed to quantify the indicated mRNAs. β-actin was used to normalize the mRNA levels, and the fold change is expressed relative to either untreated control or negative control siRNA treated cells (value set to 1). The error bars represent SD, Bonferroni test was used to calculate the significance. *P≤0.05, **P≤0.01, ***P≤0.001 and ns is non-significant i.e., P>0.05.

**Figure 2:**
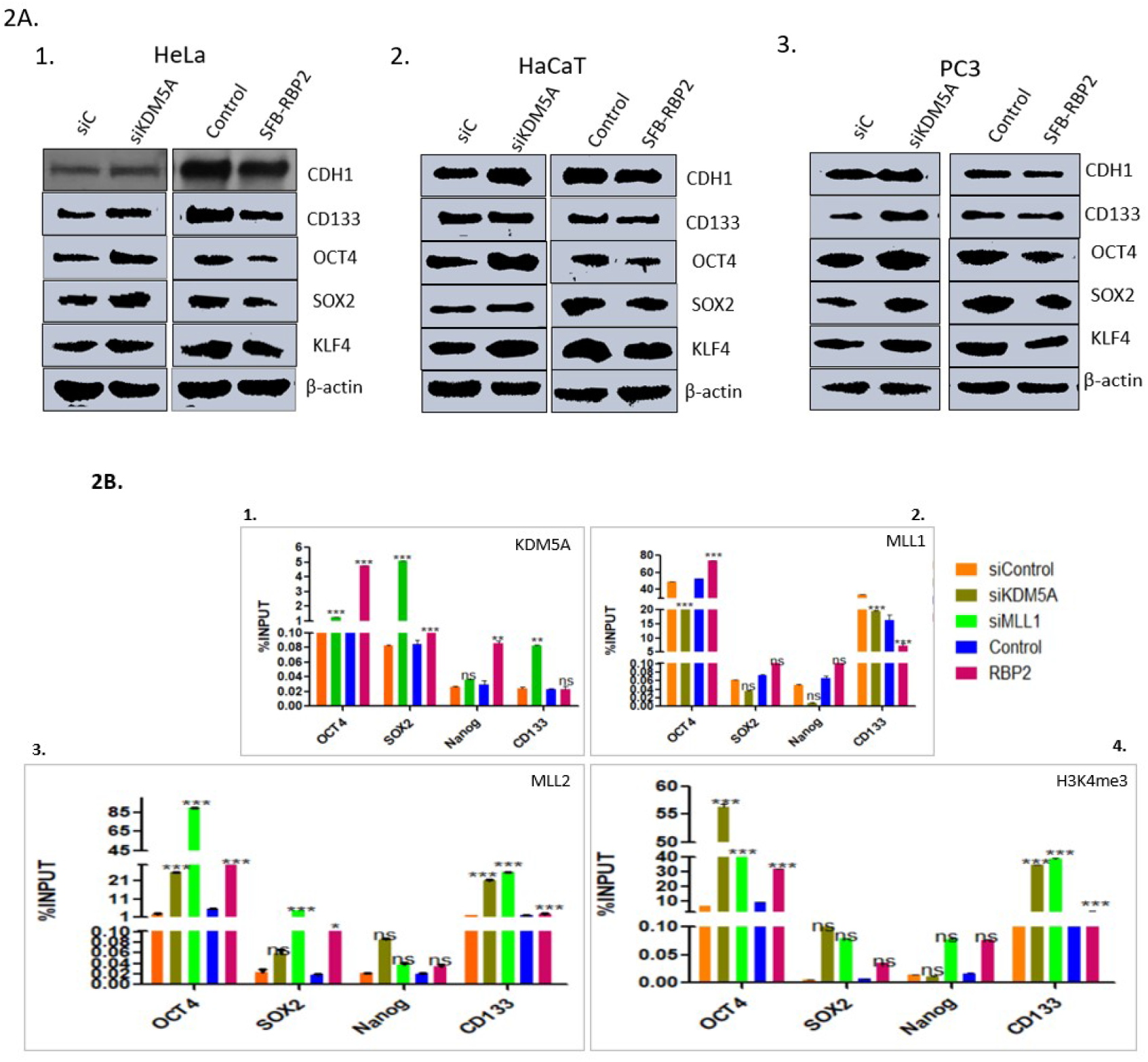
KDM5A regulates E-cadherin and stemness and pluripotency related markers directly in a demethylase dependent manner. **(A)**. Western blotting showing expression of E-cadherin, stemness (CD133) and pluripotency (OCT4, SOX2 and KLF4) markers in HeLa (1), HaCaT (2) and PC3 (3) cell lines following treatment of cells with siRNA against KDM5A and KDM5A overexpression conditions. Specific antibodies used to probe the blots are indicated to the left of each panel. **(B)**. In HaCaT cells: ChIP analysis of the promoters of stemness (CD133) and stemness (OCT4, SOX2, Nanog) markers following siRNA against KDM5A and MLL1 and overexpressing KDM5A, to study the occupancy of KDM5A (1), MLL1 (2), MLL2 (3) and H3K4me3 (4) using the indicated antibodies. IgG ChIP data is provided in supplementary figure S2. The error bars represent SD, Bonferroni test was used to calculate the significance. *P≤0.05, **P≤0.01, ***P≤0.001 and ns is non-significant i.e., P>0.05. [Please note panel 2 for MLL1 occupancy regulated by KDM5A knockdown and overexpression].

**Figure 3:**
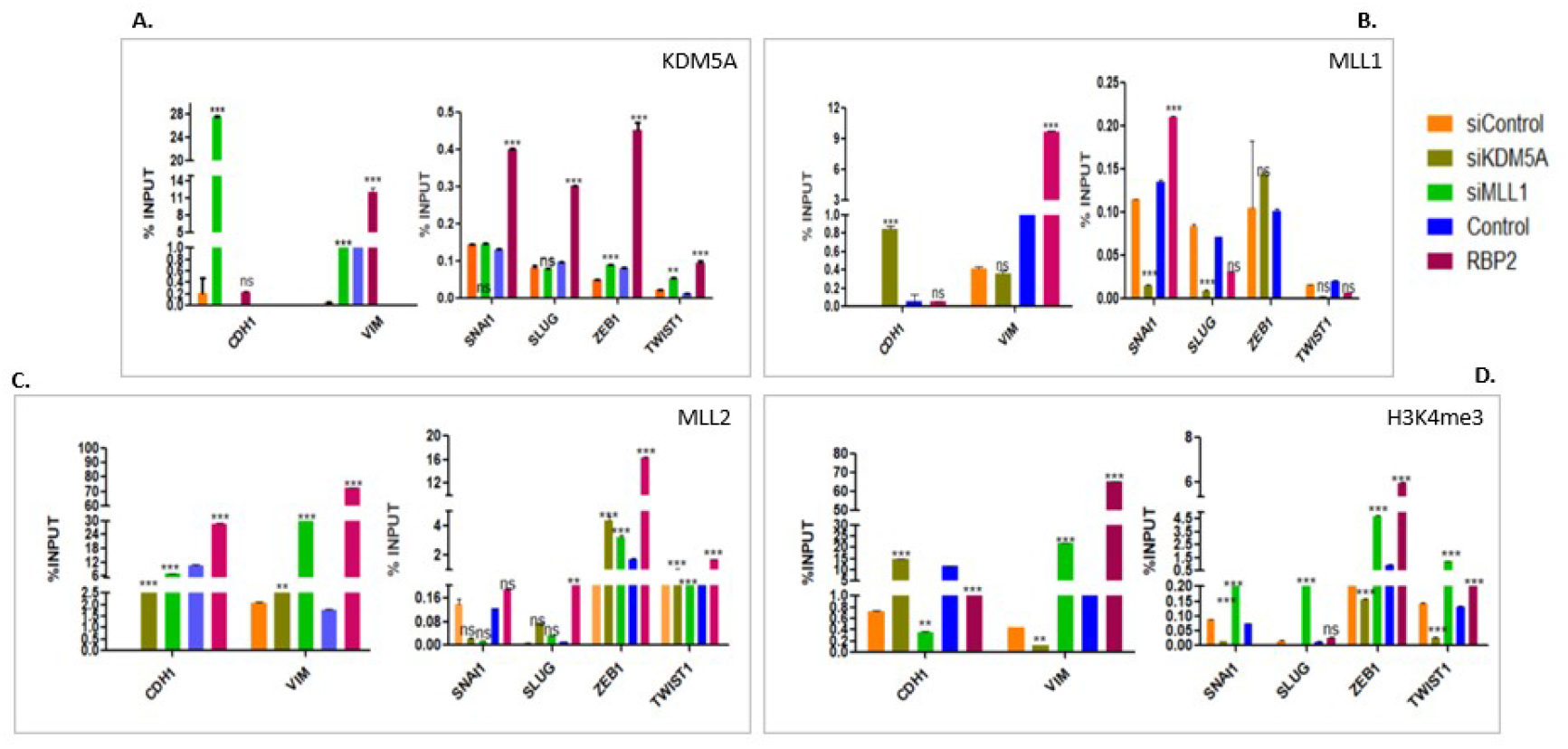
ChIP analysis of EMT markers in HaCaT cell line using siRNA against KDM5A and MLL1 and overexpressing KDM5A. Antibodies used for immunoprecipitation are indicated at the top-right corner of each panel. IgG ChIP data is provided in supplementary figure S4(IgG). The error bars represent SD, Bonferroni test was used to calculate the significance. *P≤0.05, **P≤0.01, ***P≤0.001 and ns is non-significant i.e., P>0.05. [Please note that KDM5A overexpression led to enhanced MLL2 occupancy of most the promoters studied here – Panel C].

Due to the paradoxical observations of the role of KDM5A in pluripotency using human or mouse stem cells (21,22) focused was laid on unravelling the mechanism of KDM5A mediated gene regulation on pluripotency and stemness related factors – OCT4, SOX2, NANOG (at transcript level), KLF4 and CD133 using the above-mentioned cell lines and experimental conditions. In our study using either immortalized normal cell line (HaCaT) or cancerous cell lines (HeLa and PC3), from transcript (figure 1D-F) and protein analysis (figure 2A), it turns out that KDM5A regulated all the genes under our study in a mechanism similar to E-cadherin.

To validate the data at molecular level, ChIP was performed on OCT4, SOX2, NANOG and CD133 promoters in HaCaT cell line and it was observed that KDM5A knockdown led to decreased MLL1 (figure 2B.2) occupancy (on all the 4 promoters tested) but increased MLL2 and H3K4me3 (figure 2B.3&4). It was noted that decrease in MLL1 occupancy did not affect the expression of these genes as MLL2 is the prime methyltransferase at bivalent promoters (8). In KDM5A overexpression, both MLL1 and MLL2 occupancy (on 3 of the 4 promoters tested) was enhanced (figure 2B.2&3) and MLL1 knockdown enhanced KDM5A and MLL2 (figure 2B.1&3). Irrespective of the experimental conditions, H3K4me3 remained high on all the promoters compared to the controls used (figure 2B.4), which could be justified by the fact that most of the pluripotency gene promoters are bivalent in nature, i.e., presence of H3K4me3 primes them to transcriptionally accessible state, but removal of H3K27me3 is required to transcribe them (30). Although KDM5A overexpression enhanced both MLL1 and MLL2 occupancy, it was speculated that the downregulation of these genes was probably due to co-occupancy of PRC2 components, which were earlier reported to interact with KDM5A (31). Enhanced KDM5A occupancy in MLL1 knockdown clarifies the mechanism behind regulating pluripotency genes (figure 2B.1).

### KDM5A promotes EMT in a demethylase independent mechanism

While performing KDM5A overexpression, prominent morphological changes in cells which displayed distinctive mesenchymal phenotype compared to untreated controls was observed (supplementary figure S3) which led us to assume that KDM5A overexpression alone was sufficient to induce complete EMT. As described earlier, EMT is a multistage process and in the previous section, it was presented that the mechanism of E-cadherin regulation by KDM5A is direct, i.e., its demethylase activity dependent. But, from our quantitative-PCR analysis, it turns out that KDM5A overexpression somehow enhanced mesenchymal markers expression (figure 1A-C), which doesn’t relate to the fact that KDM5A is a transcriptional co-repressor as it demethylates an active chromatin mark. The upregulation of mesenchymal markers was not MLL1 dependent as our ChIP and quantitative-PCR data reveal that MLL1 knockdown also upregulated these markers (figure 1A-B) and that KDM5A overexpression did not enhance MLL1 on 3 (Slug, Zeb1 and Twist1) of the 5 gene promoters under study (figure 3B). Further, this data showed that H3K4me3 was enhanced on all the mesenchymal promoters in KDM5A overexpression and MLL1 knockdown conditions (figure 3D), and this cannot to attributed to MLL2 occupancy which was observed to be consistently enhanced in KDM5A knockdown conditions as well (figure 3C) but couldn’t upregulate the expression (at transcript level – figure 1A-C) or increase H3K4me3 in KDM5A knockdown conditions (figure 3D).

Taken together, these observations implicate that it wasn’t MLLs that regulated mesenchymal markers expression, but instead the occupancy of KDM5A was critical and that either presence or absence of MLL1/2 did not affect their regulation. To address the question of how KDM5A, being a transcriptional repressor, activated the mesenchymal markers, previous reports on activator function of KDM5A were scrutinized and one of the mechanism by which KDM5A was reported to act as an activator was by increasing promoter acetylation by inhibiting HDAC activity (32) and from previous studies, it was also understood that Snail1, Slug and Zeb1 activation is enhanced by p300/CBP mediated acetylation of their promoter regions (33,34) and the use of p300 inhibitor (A-485) in breast cancer cell line JIMT1 led to reduced H3K18ac levels and an associated decrease in the expression of genes promoting EMT (ITGA6 and SIX2) along with reduced mesenchymal marker expression (TWIST, SNAIL, SLUG) (35). Along this line, the use of HDAC inhibitors (TSA/VA) in some cancers induced mesenchymal transition by increasing the expression of mesenchymal markers (2). Keeping these reported observations in mind, HDAC1 and H3K18ac occupancy on the mesenchymal marker promoters in KDM5A overexpression conditions was tested. Though H3K18ac occupancy was enhanced, it could be observed that HDAC1 also showed increased occupancy (figure 4). Further, to clarify on HDAC enzyme activity in KDM5A overexpression, HDAC activity assay was performed using HaCaT cell line, and it was noted that KDM5A overexpression reduced HDAC activity (supplementary figure S6). This concluded that the KDM5A mediated activation of mesenchymal markers was demethylase activity independent and was due to KDM5A mediated inhibition of HDAC activity resulting in enhanced H3K18ac.

**Figure 4:**
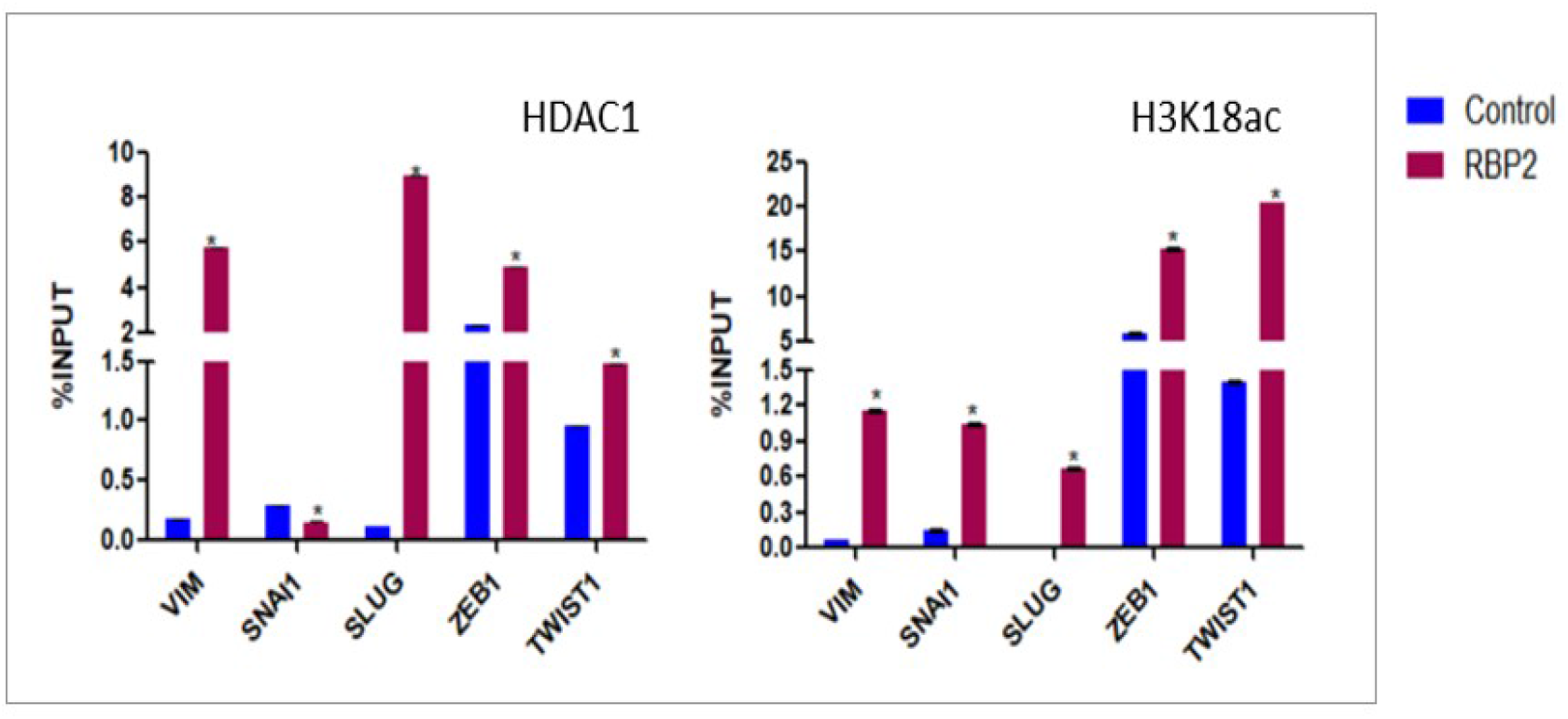
HDAC1 and H3K18ac occupancy on mesenchymal promoters following KDM5A overexpression in HaCaT cell line. The overexpression of KDM5A enhances the occupancy of both HDAC1 and H3K18ac on mesenchymal promoters and increase in the presence of acylation led to their upregulation. IgG control is represented in supplementary figure S5 (IgG). Student ‘t’ test was used to calculate significance Error bars represent SD, *P≤0.05.

All the above findings suggested that KDM5A prominently regulated the induction of EMT, which prompted us to explore the signalling pathway(s) involved in regulating cell motility. Therefore, pathway mapping using KEGG database was performed and we understood that most of the ligands that respond to tactile stimulus converge into and depend on effectors of either RTK or ITG-FAK signalling (KEGG pathway ID – hsa04810), thus MAPK and FAK signalling were targeted for preliminary investigation and it was found that KDM5A expression was not affected by MAPK pathway but inhibition of FAK affected the expression of KDM5A.

### FAK inhibition mediated attenuation of KDM5A could not alleviate EMT

As FAK is observed to be overexpressed in many cancers and FAK mediated signalling plays a pivotal role in EMT (36), we sought to explore the mechanism of FAK mediated induction of EMT in an epigenetic context. Using IC30 concentration determined from MTT assay (supplementary figure S7), HeLa and HaCaT cell lines were treated with an FAK inhibitor (PF-573228) to test its effects on E-cadherin, KDM5A and MLL2 expression. Earlier studies provided evidence to FAK signalling mediated EMT through transcriptional regulation of mesenchymal markers and by E-cadherin downregulation or delocalization and using small molecule FAK inhibitors inhibited EMT (37), but in this study, blocking FAK did not upregulate E-cadherin expression (figure 5) or inhibit EMT. This finding was unexpected as FAK inhibition reduced KDM5A and MLL2 expression (figure5), an in-vivo scenario reflecting KDM5A knockdown conditions. From our earlier experiments which revealed that KDM5A knockdown led to increase in E-cadherin expression (figure 2A), but FAK mediated decrease in KDM5A could not induce E-cadherin expression. As treatment with PF-573228 (at IC30) led to high cell death, a lower concentration of the drug was also tested in HaCaT cell line, to check if a concentration dependent effect exists, but couldn’t detect any drastic/contradictory changes in the pattern of regulation as low concentration either had equal or marginally higher E-cadherin expression and reduced KDM5A expression and higher concentration exhibited reduction in expression of both the proteins (supplementary figure S8) as seen earlier (figure 5).

**Figure 5:**
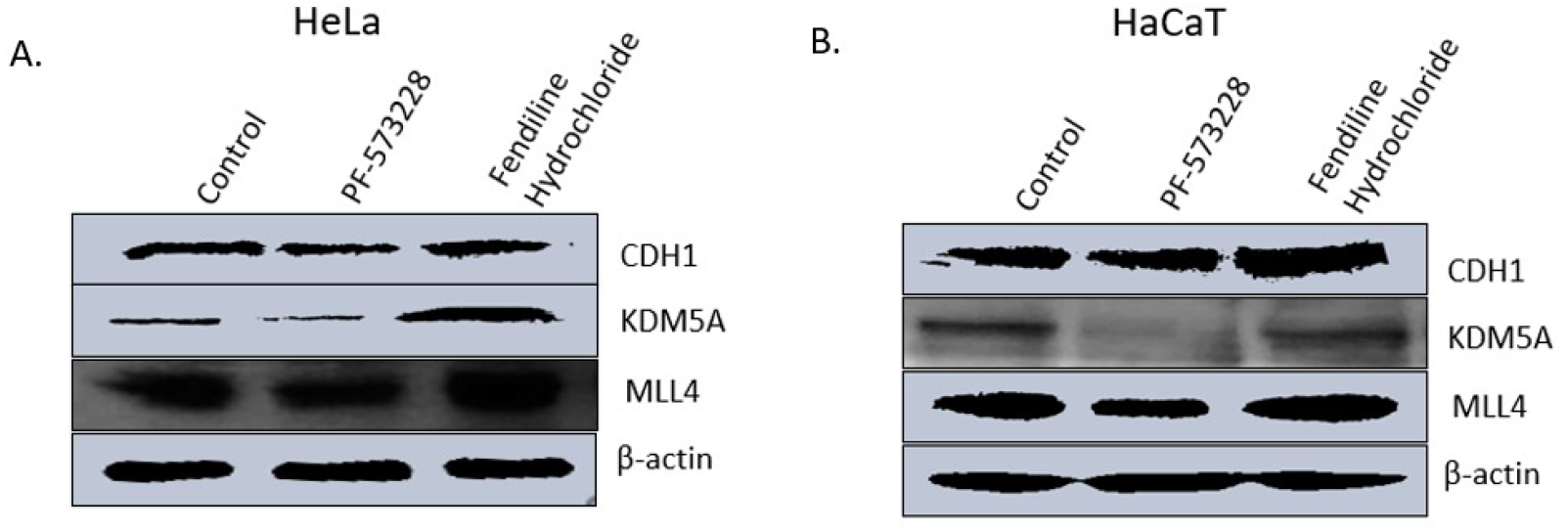
The effects of blocking FAK and calcium signalling pathway on the expression of E-cadherin, KDM5A and MLL2 in HeLa (Panel A) and HaCaT (panel B) cell lines. Protein from cells treated with the following drugs were subjected to immunoblot and probed with the antibodies indicated on right side of each panel. A pattern of co-expression was observed among all the three proteins tested here.

When similar experiments were performed in other cell lines (HCT15 and A498, our unpublished findings), which showed increase in E-cadherin, KDM5A and MLL2, contradicting our observations from HeLa and HaCaT cell lines. These observations in HCT15 and A498 can be theoretically validated as FAK inhibition reduces EMT and enhances E-cadherin, but still does not explain the KDM5A and MLL2 regulation over E-cadherin, as blocking FAK pathway in these cell lines led to increase in KDM5A – that mimic the experimental conditions of KDM5A overexpression, but instead of downregulating E-cadherin, enhanced expression was observed.

To decode the mechanism associated with FAK inhibition mediated E-cadherin expression and EMT, ChIP experiments were performed in HaCaT cell line. Interestingly, FAK inhibition led to reduced KDM5A and MLL2 occupancy (figure 6 A&C) associated with reduced H3K4me3 (figure 6D) on E-cadherin promoter. From this and our previous experiments (figure 3), it was understood that both MLL1 and MLL2 are required for inducing E-cadherin expression and reduction in the occupancy of either MLL1 (in case of knockdown) or MLL2 (in case of PF-573228 treatment) decreased E-cadherin expression. In line with downregulating E-cadherin, KDM5A occupancy on mesenchymal markers was seen to be enhanced (figure 6A), concluding that FAK mediated downregulation of KDM5A didn’t alleviate EMT induction.

**Figure 6:**
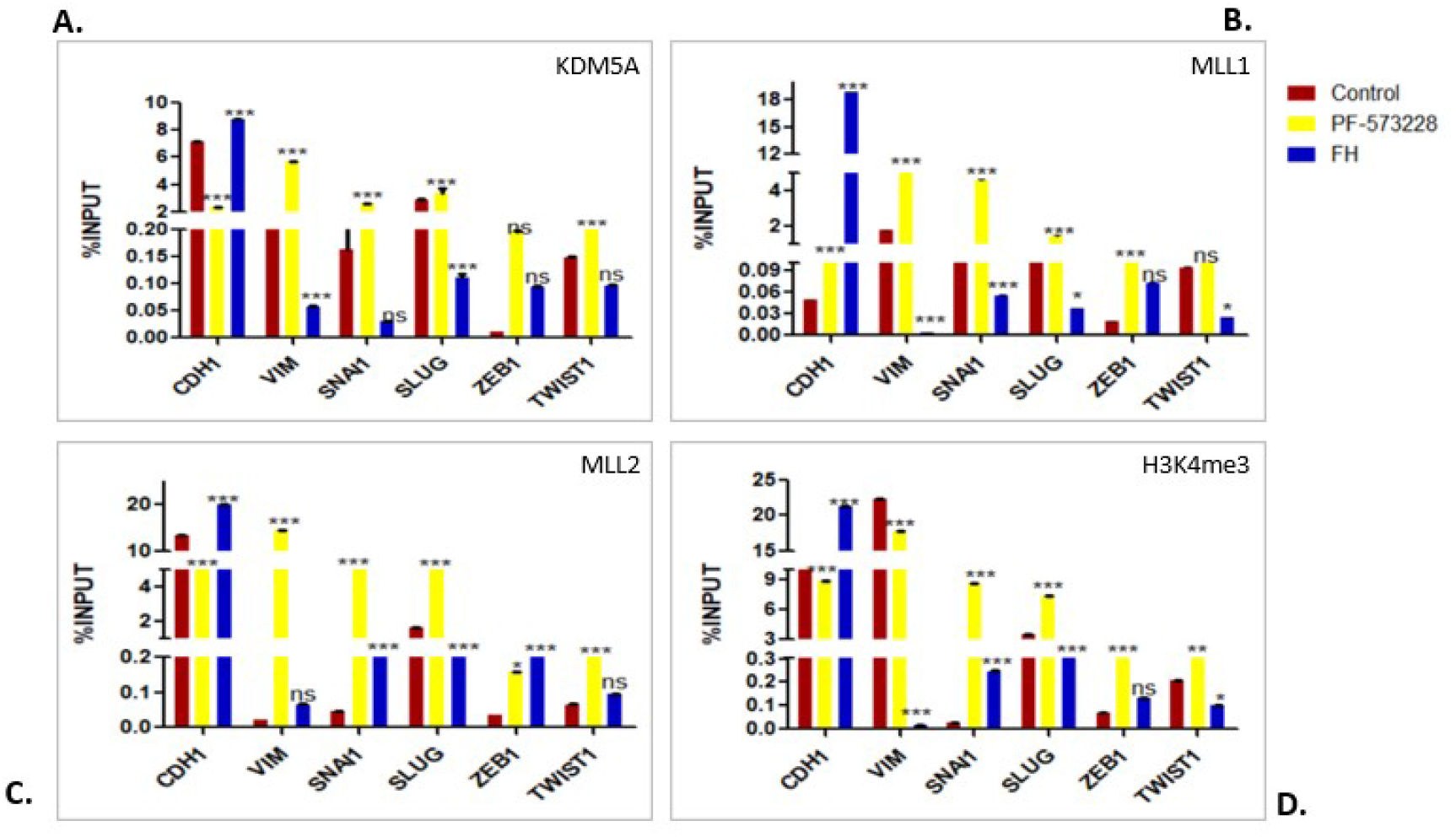
ChIP analysis following treatment of HaCaT cells with FAK (PF-573228) and calcium signalling (fendiline hydrochloride -FH) inhibitors on mesenchymal marker promoters using antibodies as indicated on the top right corner of each panel. IgG occupancy of the following experiment is presented in supplementary figure S10. The error bars represent SD, Bonferroni test was used to calculate the significance. *P≤0.05, **P≤0.01, ***P≤0.001 and ns is non-significant i.e., P>0.05.

As FAK signalling is reported to enhance pluripotency (20,38), we further examined if FAK inhibition affected the stemness marker CD133, and observed that blocking FAK pathway did reduce stemness (supplementary figure S9), in a mechanism similar to the regulation of E-cadherin by KDM5A and MLL2, i.e., reduced MLL2 occupancy mediated downregulation was observed, adding strength to the existing literature that MLL2 is the methyltransferase at bivalent promoters.

### Elevated KDM5A levels following calcium signalling inhibition did not repress E-cadherin expression

Intracellular calcium is a crucial regulator of EMT which acts via modulating FA turnover during metastasis and by stabilizing the filopodia formation (39,13). Similar to FAK inhibition, calcium signalling (refer to supplementary figure S7 for MTT assay) was also inhibited using fendiline hydrochloride, to study the signalling-epigenetic interplay. As calcium signalling enhances EMT, consistent with previous reports (40,41), our study also shows that chelating intracellular calcium levels induced E-cadherin expression (figure 5), associated with increased KDM5A and MLL2 expression. Again, like in FAK inhibition, the general trend of KDM5A mediated E-cadherin regulation was not observed. Instead, enhanced KDM5A led to increased E-cadherin expression. Our ChIP experiments provided evidence that enhanced MLL2 and H3K4me3 occupancy (figure 6 C&D) associated with increased KDM5A (figure 6A) on E-cadherin promoter. On promoters of genes involved in mesenchymal transition, KDM5A occupancy reduced (figure 6A), thereby downregulating their expression.

### FAK signalling is required for KDM5A enzymatic activity

As mentioned earlier, inhibition of FAK signalling (in HCT15 and A498) or chelation of calcium signalling (in HeLa and HaCaT) enhanced KDM5A expression, and from KDM5A overexpression studies, we know that presence of KDM5A downregulates E-cadherin irrespective of MLLs occupancy on the promoter, but in both the drugs treatments, enhanced KDM5A occupancy on E-cadherin promoter did not downregulate its expression, leading us to hypothesize that the enhanced KDM5A protein is catalytically inactive and only mediated MLL2 recruitment to the E-cadherin promoter. To confirm the same, we exploited the PF-573228 treatment conditions (which downregulated the endogenous KDM5A protein), and overexpressed KDM5A transiently to attain conditions that mimic HCT15 and A498 treatment. Interestingly, transient overexpression of KDM5A in FAK inhibited-HeLa and HaCaT cell lines enhanced E-cadherin expression (figure 7A).

**Figure 7:**
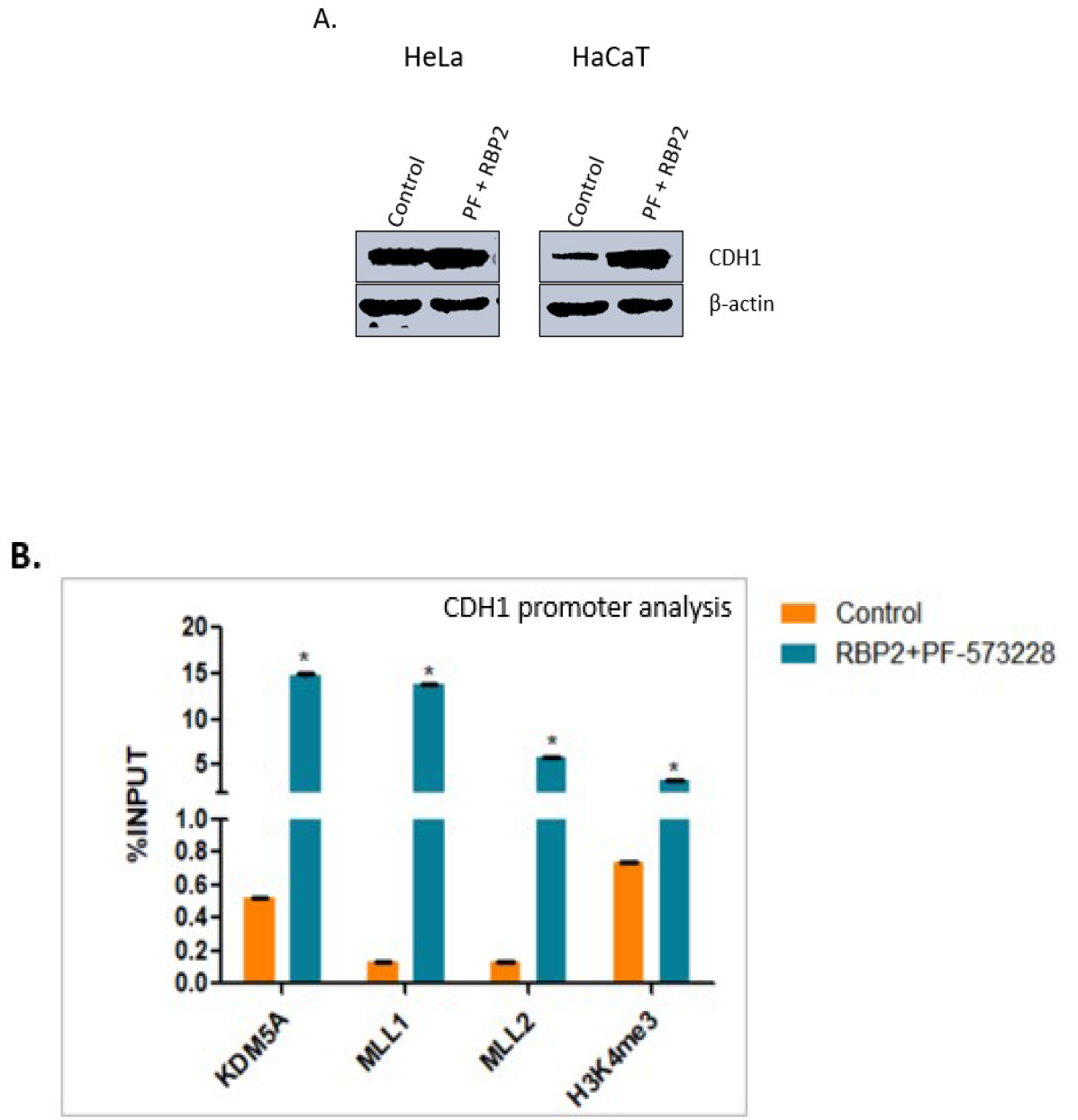
**(A)**. Transient overexpression of KDM5A in FAK signalling inhibited (by PF-573228) HeLa and HaCaT cells led to increase in E-cadherin expression. Cells treated with both PF-573228 and RBP2 construct were subjected to immunoblot and probed with E-cadherin antibody. Antibodies used are indicated to the right of the panel. **(B)**. Similarly treated HaCaT cells were subjected to ChIP analysis to study E-Cadherin promoter using antibodies indicated. Transient overexpression of RBP2 in PF-573228 treated cells led to enhanced occupancy of both MLL1 and MLL2 compared to the control. IgG is represented in supplementary figure S11. Student ‘t’ test was used to calculate significance Error bars represent SD, *P≤0.05.

Motivated by this interesting observation, we further tested if this enhanced KDM5A occupancy tethered MLL2 to E-cadherin promoter and increased H3K4me3 leading to its upregulation, ChIP experiments were designed in FAK inhibited-transient KDM5A overexpressing HaCaT cells, and as per our expectation, both KDM5A and MLL2 occupancy increased in these treatment conditions (figure 7B), which was associated with increased H3K4me3 levels (figure 7B) in comparison to their reduced occupancy in FAK-inhibition treatment alone (figure 6A,C-D).

### KDM5A enhances the occupancy of MLL1 or MLL2 on gene promoters

KDM5A and MLL2 co-expression (figure 5) was observed following drug treatments, similar results were obtained in HCT15 and A498 (our unpublished observation) cell lines as well. Throughout our ChIP experiments, it was found that KDM5A co-occupied with either of the MLLs. On the bivalent promoters (i.e., pluripotency and stemness related gene promoters) KDM5A knockdown reduced MLL1 and its overexpression enhanced MLL1 occupancy on all the promoters studied (figure 2B.2). On EMT marker promoters, KDM5A overexpression enhanced MLL2 occupancy on E-cadherin and mesenchymal promoters (figure 3C) along with co-occupancy of KDM5A and MLL2 observed in both the drug treatments (figure 6 A&C). KDM5A mediated recruitment of MLL2 to E-cadherin promoter was confirmed when KDM5A was transiently overexpressed in PF-573228 treated HaCaT cells which led to enhanced occupancy of MLL2 (figure 7B) and upregulated E-cadherin.

### Physical association of KDM5A with MLL1 and MLL2

Theoretically, as proteins that co-express and co-occupy were reported to interact with each other, hence we wanted to see if KDM5A interacts with MLL1 and MLL2 and aid in their recruitment to target promoters. To examine the same, co-immunoprecipitation experiments were performed with endogenous proteins in U87-MG and HaCaT cell lines. Using either MLL1, MLL2 and KDM5A antibodies with A/G beads, we immunoprecipitated whole cell lysates of U87MG and HaCaT cell lines, and the samples were subjected to western blotting. The co-immunoprecipitation experiments showed that KDM5A interacted with both MLL1 and MLL2 (in U87MG - figure 8B) and immunoprecipitation using KDM5A antibody also detected MLL2 interaction in U87MG (figure – 8A) and in HaCaT (supplementary figure S12).

**Figure 8:**
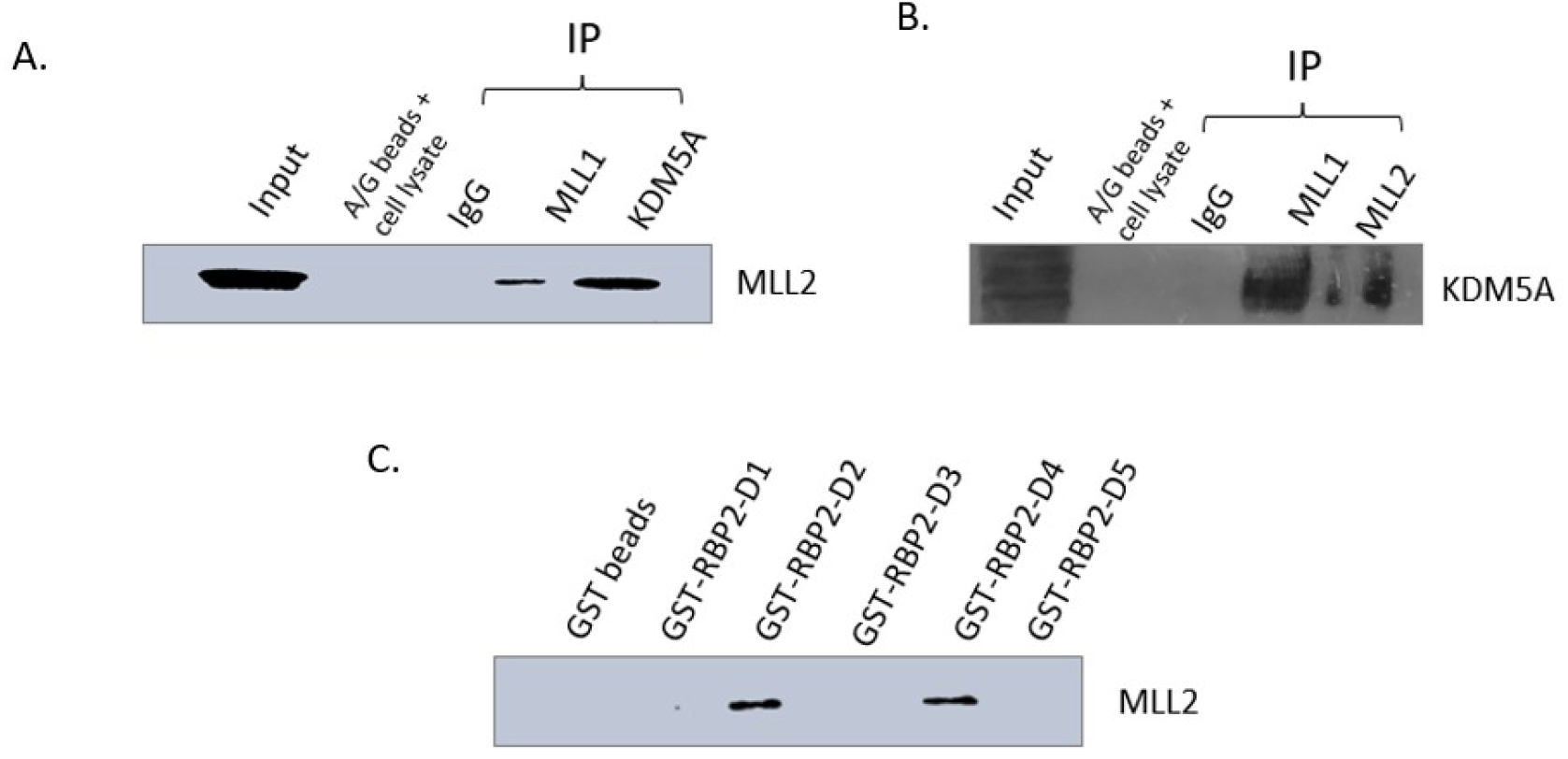
KDM5A interacts with MLL1 and MLL2. **(A)**. MLL2 associates with endogenous MLL1 and KDM5A proteins when immunoprecipitated using respective antibodies and immunoblot is probed with MLL2. **(B)**. KDM5A also interacted with endogenous MLL1 and MLL2 proteins in co-IP experiments using U87MG cell extracts where IP was performed usingMLL1 and MLL2 antibodies and the blot is probed with KDM5A. **(C)**. Bacterially expressed GST-tagged RBP2 deletions (D1-D5) were used for pulldown to locate MLL2 interaction. It was observed that D2 and D4 interacted with endogenous MLL2 protein from U87MG cells. Nomenclature of deletion constructs as per the original publication (Zargar et al., 2018): D1 – contains JmjN and ARID domain; D2 – PHD1-JmjC-ZF; D3 – PLU-1 like; D4 – PHD2; and D5 – PHD3.

### MLL2 binds to the catalytic domain and PHD2 domain of KDM5A

Further, to map the exact region of KDM5A that interacted with MLL2, pull-down experiments were setup with GST-fusion fragments of KDM5A which were overexpressed in BL21-DE3, purified using glutathione-GST beads, and used to pulldown endogenous protein from whole cell extracts of U87MG and HaCaT cells. Five deletion constructs of KDM5Awere used (D1-D5; where D1 denotes JmjN and ARID domain; D2 – PHD1-JmjC-ZF; D3 – PLU-1 like; D4 – PHD2; and D5 – PHD3 - borrowed from Dr. Shweta Tyagi (Zargar et al., (2018)). Both fragment D2 and D4 interacted with MLL2, in both the cell lines (figure 8C and supplementary figure S12). Fragment D2 contains PHD1-JmjC-ZF domains of KDM5A and fragment D4 contains PHD2 domain. This interaction mapping data implicates that MLL2 uses PHD2 to tether to KDM5A and might block KDM5A enzymatic activity by binding to the PHD1-JmjC region.

## Discussion

Gene expression is orchestrated by interplay between signalling pathways and epigenetic machinery. Components of signalling pathways can either interact or post-translationally modulate the activity of enzymes and co-factors of epigenetic machinery and epigenetic modifiers in turn regulate the expression and activity of many proteins in a cell (42). H3K4me3 is a transcriptionally active mark, laid down by SET domain containing methyltransferases like MLL1 and MLL2 and erased by JmjC family of demethylases like KDM5A. Many signalling pathways and epigenetic modulators control the process of EMT, that involves downregulation of epithelial markers and enhancement of mesenchymal marker expression, and the exact mechanism of how KDM5A and MLLs control these two sets of genes in not yet known. Also, pluripotency related bivalent promoters are generally controlled by H3K4me3 and H3K27me3; and previous studies of KDM5A’s regulatory mechanism at bivalent promoters is contradictory.

Here, in this study, following our preliminary transcript and protein quantification experiments in 3 cell lines using knockdown (KDM5A and MLL1) and overexpression (KDM5A) approaches, it was understood that all of the stemness and pluripotency genes and E-cadherin are direct targets of KDM5A and MLL1, meaning that at these promoters, both these epigenetic modifiers function as classic demethylase or methyltransferase respectively. But at mesenchymal promoters, it was noted that occupancy of KDM5A somehow induced these genes, and absence of MLL1 didn’t render them inactive, concluding that KDM5A activates EMT and MLL1 is dispensable (from our MLL1 knockdown studies) for induction of this process. Our ChIP experiments validated the fact that E-cadherin is under the control of MLL1 and KDM5A and mesenchymal markers are activated by the presence of KDM5A. In case of the bivalent promoters, KDM5A renders them inactive and MLL2 is more important for their induction. Interestingly, we observed co-occupancy of KDM5A with MLL1 on pluripotency promoters and with MLL2 on EMT related promoters. As for the KDM5A-mediated activation of EMT promoters, it was found that KDM5A-mediated inhibition of HDAC activity on these promoters led to enhanced H3K18ac thereby upregulating the mesenchymal markers in the presence of KDM5A.

As FAK and calcium signalling are dysregulated in many cancers and both these pathways critically influence EMT and pluripotency (via integrin-FAK signalling), we sought to elucidate if these signalling pathways regulate KDM5A and MLL2 and thus influence gene expression. Through this work, we demonstrate that KDM5A – mediated regulation of E-cadherin promoter follows different mechanisms, as FAK inhibition mediated downregulation of KDM5A didn’t enhance cadherin or chelating calcium mediated increase in KDM5A couldn’t repress cadherin expression. As MLL2 and KDM5A were co-expressed in both the drugs treatments, ChIP was performed to study their occupancy on EMT promoters. Similar to our previous observations of KDM5A and MLL1 co-occupancy of pluripotency promoters, a depletion of MLL2 was seen whenever KDM5A occupancy reduced and also that enhanced KDM5A occupancy led to increased MLL2 on E-cadherin promoter, leading us to the conclusion that KDM5A recruited MLL2 to gene promoters which was validated by transient overexpression of KDM5A in PF-573228 treated cells that led to increased MLL2 occupancy. As the catalytically inactive or kinase-dead mutants of FAK generally localize to nucleus and nuclear FAK regulate gene expression by interacting with epigenetic modifiers like HDAC1, MBD2 and Sin3a (43), all of which interact with KDM5A (9) and increased KDM5A occupancy didn’t downregulate E-cadherin expression, it was concluded that FAK inhibition somehow rendered KDM5A inactive. We speculate that a phosphorylation dependent activation mechanism might persist between FAK and KDM5A as KDM5 family members localization and enzymatic activity was earlier reported to be regulated by phosphorylation through AKT/CDKs (44,45). The differential expression profile of KDM5A-MLL2 in HeLa-HaCaT cell lines (this study) compared to HCT15-A498 (our unpublished data) could be attributed to FAK localization in these cell lines as in colon cancer, phosphorylated FAK localized to nucleus compared to ovarian cancer which exhibited both cytosolic and nuclear distribution (43).

From our protein expression and chromatin immunoprecipitation analysis, we observed co-occupancy of KDM5A with either MLL1 or MLL2, leading to hypothesize if there was any possible interaction between these antagonistic players? This study confirms that KDM5A indeed interacts with both MLL1 and MLL2 in a context dependent manner, and for the first time, we assign functionality to the PHD2 domain of KDM5A as MLL2 interacted with the PHD2 and catalytic region (PHD1-JmjC) of KDM5A. Though this interact requires structural validation, functionally, it was understood that MLL2 might use PDH2 domain as an anchoring junction and block the catalytic domain of KDM5A.

Taken together, different modes of gene regulation were elucidated in this study -starting with the observation that presence of both MLL1 and MLL2 and absence of enzymatically active KDM5A drives E-cadherin and bivalent promoters, but in case of bivalent genes, MLL2 is the prime methyltransferase – all of which is consistent with the existing mechanism of MLL-KDM5A enzyme activity. On mesenchymal promoters – KDM5A acts an activator by HDAC activity inhibition mediated increase in expression. Although many interactions of KDM5A with other epigenetic players are known (9), interesting was the observation that two proteins with antagonistic functions interact and regulate gene expression. From these experiments, we could picture that both KDM5A and MLL occur together at target gene promoters, and upstream signalling dictates the fate of regulation probably by adding post-translational modifications which might regulate the structural conformation, enzymatic activity and interactions between two proteins.

## Supporting information

Kirtana et al KDM5A-MLL1-2 interactions NAR

## Supplementary data statement

Supplementary data are available at NAR online

## Acknowledgements

We thank Dr. Shweta Tyagi, CDFD Hyderabad for SFB-RBP2 and GST-RBP2 deletion (D1-D5) constructs.

## Author Contributions

SKP conceptualized the project. K.R and S.M performed all the experiments, K.R designed and analysed the experiments and wrote the manuscript, and SKP edited the final manuscript.

