## Supplementary material for "Physical association of functionally antagonistic enzymes: KDM5A interacts with MLLs to regulate gene expression in a promoter specific manner facilitating EMT and pluripotency": Kirtana et al KDM5A-MLL1-2 interactions NAR

#### Supplementary information

#### Supplementary methods:

##### Cell culture conditions

HeLa and HaCaT cells were cultured in MEM and DMEM media respectively, with 10% FBS (Gibco -10270106) and 1% anti-anti. PC3 was grown in F12 media supplemented with 15% FBS, 1% anti-anti and (Gibco -15240-062) L-glutamine. For co-immunoprecipitation studies, we used U87MG cells that were maintained in complete DMEM media. During treatments, depending on cell doubling size, seeding density was calculated to attain 70% confluency within 24hrs, following transfections or drug incubations. For microscopy experiments, seeding density was reduced to avoid overcrowding.

##### siRNA transfections and pcDNA-SFB-RBP2 plasmid overexpression

In HeLa and HaCaT cell lines, both knockdown and overexpression was performed for 48hrs, but in PC3 the treatment duration was 24hrs. Transfection was attained using lipofectamine 3000 (Invitrogen L3000-15) following manufacturer's instructions. Plasmid concentration varied from 6 well to 60mm plate and we used 5 or 10ug respectively to induce KDM5A overexpression. The pcDNA-triple epitope SFB-tagged-RBP2 (KDM5A) overexpression construct was borrowed from Dr. Shweta Tyagi (26).

Efficiency of knockdown and overexpression was confirmed by immunoblotting (supplementary figure S1-A). As MLL1 is a huge protein (432KDa), and we couldn't perform immunoblotting successfully, so we analysed its knockdown efficiency by immunocytochemistry (supplementary figure S1-B).

**siRNAs used in this study:**
KDM5A siRNA: Thermo – siRNA ID – s11836 (Ambion silencer select siRNA)
Sequence: Sense – GCGAGUUUGUUGUGACAUUtt; Antisense – AAUGUCACAACAAACUCGCca
MLL1 siRNA: Sigma – siRNA ID – SASI_Hs01_00090459
Negative control: Thermo – catalog no – 4390843 (Silencer select negative control siRNA)

##### Immunocytochemistry

HeLa and HaCaT cells were seeded on a coverslip in a 6-well plate, transfected with MLL1 siRNA (30nM) for 48hrs followed by fixation using 100% methanol for 5 minutes, permeabilized with PBST on ice for 10 minutes and blocked using 1% BSA in PBST for 1hr. The cells were incubated with anti-MLL1 antibody overnight at 4$^{o}$C in a humid chamber. Following 3 washes with PBS, the cells were incubated in Alexa647 (anti-mouse – ab150119), washed thrice and images were acquired using Leica microsystems.

**Quantitative real time PCR**

To quantify the changes in gene expression profile following treatments, RT-PCR was performed. RNA isolation was performed by TRIzol (Thermo - 15596018) method, using isopropanol (himedia-MB063) for RNA precipitation and 70% alcohol (Himedia-MB228) to wash off excess salts and contaminants, and the RNA was dissolved in DEPC water. Reverse transcriptase reaction was setup to synthesize cDNA as per manufacturer's instructions (Genesure – PGK163A) with the above isolated RNA with incubation at 42°C for 60min and 70°C for 5 min using a 20uL reaction mixture. Using this cDNA (1 ug) as template, and gene specific primers (300nM) respective to different physiological processes, RT-PCR was performed using SYBR-green technology (Thermo – A25742) with the following cycling conditions 50.0°C for 1:00; 96.0°C for 6:00 and [96.0°C for 0:10; 55.0°C for 0:30; 72.0°C for 1:00] for 40 cycles. The data was analysed according to Livak's method (ddCT calculation) using either GAPDH or B-actin as control using formula adapted from (46).

**MTT assay and drug treatments**

Cell viability changes following drug treatment were determined by measuring absorbance off MTT (Himedia-TC191) in living cells. In brief, 24hrs prior to the drug administration, $10^3$ to $1.5*10^3$ cells were seeded into a 96-well plate, followed by replacing the normal media with drug dissolved media at required dilutions. The plate was then replaced into the incubator for 24hrs for the drug to be effective, followed by addition of MTT media, which was incubated for another 6hrs followed by DMSO (Himedia – AS121) mediated dissolution of the formazan crystals. A colorimetric analysis of the optical density at 570nm gave us the percentage of viable cells following treatment with different concentrations of a particular drug.

The percentage of viability was calculated as follows:

% Viability =100 * mean OD (drug)/mean OD (control)

Drugs used in this study: The FAK phosphorylation inhibitor – PF-573228 was purchased from Sigma (PZ0117-5MG) and LTCC calcium channel inhibitor – Fendiline hydrochloride was from Santacruz (SC239958). Cells were treated with these drugs foe 24hrs prior to downstream experimentation. Graphs of %viability obtained from MTT are presented in supplementary figure 4.

**Chromatin immunoprecipitation buffers:**

ChIP lysis buffer - (1% SDS, 10 mM EDTA, 50 mM Tris-HCl, pH 8, and protease inhibitors)
ChIP dilution buffer - (0.01% SDS, 1% Triton X-100, 1.2 mM EDTA, 16.7 mM Tris-HCl, pH 8, 167 mM NaCl, and protease inhibitors)
low-salt buffer- (0.1% SDS, 1% Triton X-100, 2 mM EDTA, 20 mM Tris-HCl, pH 8, 150 mM NaCl)
high-salt buffer -(0.1% SDS, 1% Triton X-100, 2 mM EDTA, 20 mM Tris-HCl, pH 8, 500 mM NaCl)
LiCl buffer (250 mM LiCl, 1% NP-40, 1% deoxycholate, 1 mM EDTA, 10 mM Tris-HCl, pH 8)
TE buffer (10 mM Tris-HCl, pH 7.6, 1 mM EDTA)
Elution buffer - (0.1 M NaHCO3, 1% SDS) (freshly made)

% INPUT was calculated as follows:

% INPUT = 100* 2x-y

(Where X is the ct value for Input adjusted for dilution factor and Y is the ct value for the IP samples (26).

**HDAC activity assay**

HDAC activity assay was performed using kit (BioVision – catalog #K331-100) according to manufacturer's instructions. In a 96 well plate, whole cell lysate (100ug) was used and diluted to 85ul with ddH2O. 10uL of HDAC assay buffer and 5uL of HDAC colorimetric substrate were added and the plate was incubated at 37°C for 1hr. Further, 10uL of lysine developer was added to each sample well and incubated 37°C for another half an hour, followed by acquisition of O.D at 405nm.

**Primer's list:**

**Primers used for real-time quantitative PCR:**

1. **CDH1**
   **F:** CGAGAGCTACACGTTCACGG
   **R:** GGGTGTCGAGGGAAAAATAGG
2. **VIM**
   **F:** GGTTCAGGTTTCATTCATGCCT
   **R:** AGTTGGCTGTGTGTACTGCT
3. **Snai1**
   **F:** CCTGTCTGCGTGGGTTTTTG
   **R:** ACCTGGGGGTGGATTATTGC
4. **Slug**
   **F:** CTCACTCGCCCCAAAGATGA
   **R:** CCTCCCTTTTCTTTCCCAGTG
5. **Zeb1**
   **F:** CCCTCCTGCACCAAGAAAGAT
   **R:** TGAGGGGCTTTCCCTTTAGA
6. **Twist1**
   **F:** GCACTGTTCTTATCACCACCAC
   **R:** ACGGGTAAGGACCGTTTTG
7. **Oct4**
   **F:** AGCAAAACCCGGAGGAGT
   **R:** CCACATCGGCCTGTGTATATC
8. **Sox2**
   **F:** GGAAATGGAGGGGTGCAAAAGAGG
   **R:** TTGCGTGAGTGTGGATGGGATTGGTG
9. **Nanog**
   **F:** TCCTCCTCTTCCTCTATACTAAC
   **R:** CCCACAATCACAGGCATAG
10. **GAPDH**
    **F:** TGTTGCCATCAATGACCCCTTC
    **R:** CTCCACGACGTACTCAGCGC
11. **B-actin**
    **F:** CTGGAACGGTGAAGGTGACA
    **R:** AAGGGACTTCCTGTAACAACGCA

**Primers used for ChIP study:**

1. **CDH1**
   **F:** GTTCAGACTCCAGCCCG
   **R:** CTGCGGCTCCAAGGG
2. **Vimentin**
   **F:** CTAACCAACGACAAAGCCC
   **R:** AGCTACTTGCATGGGCG
3. **Snai1**
   **F:** CTCTTTCCTCGTCAGGAAGC
   **R:** AACGCACCTGGATTAGAGTC
4. **Slug**
   **F:** TTACGAACTGAGCCCGTTT
   **R:** AGGGAGGAGCTGAAATCTGA
5. **Zeb1**
   **F:** TACCTTTCCAACTCCGACAG
   **R:** AAGTTTTCCTCAGGTGTGGT
6. **Twist1**
   **F:** AAATCGAGGTGGACTGGGAA
   **R:** CCGGAGACCTAGGTAAGGAC
7. **CD133**
   **F:** CACAGTGTTGGCCCATTTC
   **R:** AAGGAAACTCTAGGGATGGC
8. **OCT4**
   **F:** ACCACCTTAGTGGAAGAGGG
   **R:** AAATGCCTGGTTGGAATGGA
9. **SOX2**
   **F:** AATACTGTGCTCAGCCAAGAAA
   **R:** ATTAGCACATGATGCTGGAC
10. **Nanog**
    **F:** GCATGGCAAATCACGATGAG
    **R:** CTGCCCAGTAACATCCACAA

**Antibodies used in this study:**

The antibodies against KDM5A (ab70892), E-cadherin (ab40772), OCT4 (ab18976), SOX2 (ab97959), HDAC1 (ab7028), H3K18ac (ab40888), B-actin (ab8227) - all of which were procured from Abcam, H3K4me3 (Invitrogen MA5-11199) and CD133 (NBP2-44247) purchased from NOVUS, are raised in rabbit. KMT2a (ab32400-100), KMT2b (ab56770), KLF4 (ab130243) – procured from Abcam, were mouse raised antibodies. IgG (M8695) from sigma was used in ChIP and Co-IP studies.

**Supplementary Results:**

**Figure legends**

**S1**: **(A)**. Western blotting confirming the efficiency if knockdown of KDM5A using siRNA in HeLa (A), HaCaT (B) and PC3 (C) cell lines. KDM5A overexpression using SFB-RBP2 construct was confirmed in HeLa (D), HaCaT (E), and PC3 (F) cell lines following transfection for 48hrs. **(B).** MLL1 knockdown using siRNA was confirmed by immunoblotting using MLL1 antibody, followed by staining with Alexa647.

**S2**: IgG occupancy on stemness (CD133) and pluripotency (OCT4, SOX2 and Nanog) markers in a ChIP experiment performed using HaCaT cell line with KDM5A and MLL1 siRNA and KDM5A overexpression construct, along with negative control siRNA or untreated control cells used as control. Error bars represent SD.

**S3**: Morphological changes observed in HeLa cells following KDM5A overexpression using SFB-RBP2 construct following 2 days of transfection. Induction of fibroblast like mesenchymal phenotype was observed following both 24hr and 48hr treatment with the overexpression construct when compared to untreated control cells.

**S4**: ChIP using IgG antibody, in HaCaT cell line following either KDM5A or MLL1 knockdown and KDM5A overexpression. The occupancy is ranked on promoters of EMT markers. Error bars represent SD.

**S5**: Following overexpression of KDM5A, occupancy of IgG was scored on promoters of mesenchymal markers. Error bars represent SD.

**S6**: The graph represents the colorimetric estimation of HDAC enzymatic activity following KDM5A overexpression in HaCaT cell line, measured as absorbance at 405nm. Error bars represent SD.

**S7**: The graph shows the cellular growth changes in HeLa and HaCaT cell lines following administration of PF-573228 and fendiline hydrochloride as assessed by MTT assay.

**S8**: Western blotting showing expression of E-cadherin and KDM5A following treatment of HaCaT cells with lower (IC10) and higher (IC30) doses PF-573228 drug that blocks FAK signalling. Decrease in the expression of both proteins was observed when higher drug concentration was used, but at lower concentrations, there was either a slight decrease or no change in the expression.

**S9**: Similar to E-cadherin expression, the expression of CD133 decreased in both cell lines (HeLa and HaCaT) following treatment with PF-573228, indicating a similar mechanism of regulation.

**S10**: Chromatin immunoprecipitation using IgG on EMT marker promoters following treatment of HaCaT cells with FAK (PF-573228) and calcium signalling (FH) blockers. Error bars represent SD.

**S11**: IgG occupancy on E-cadherin promoter following treatment of HaCaT cells with PF-537228 and KDM5A overexpression construct. Error bars represent SD.

**S12**: KDM5A and MLL2 interaction analysed using HaCaT cell extracts, **(A)**. KDM5A interacts with MLL2 when endogenous protein was subjected to immunoprecipitation using KDM5A antibody. **(B)**. GST-tagged RBP2 deletions were expressed in bacteria and purified using glutathione-agarose beads, and purified bead bound KDM5A deletion fragments were incubated with whole cell extracts of HaCaT to detect interaction with endogenous MLL2 protein, and we observed that both D2 and D4 could associate with MLL2.

#### Supplementary figures

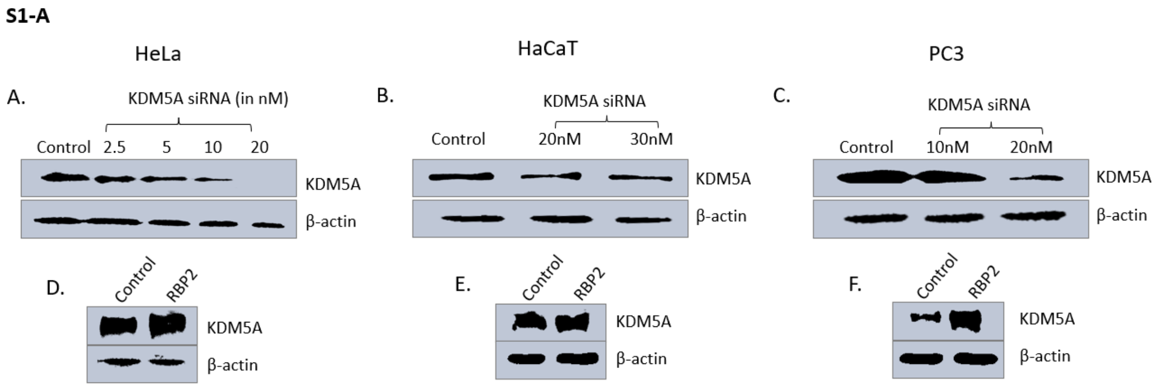


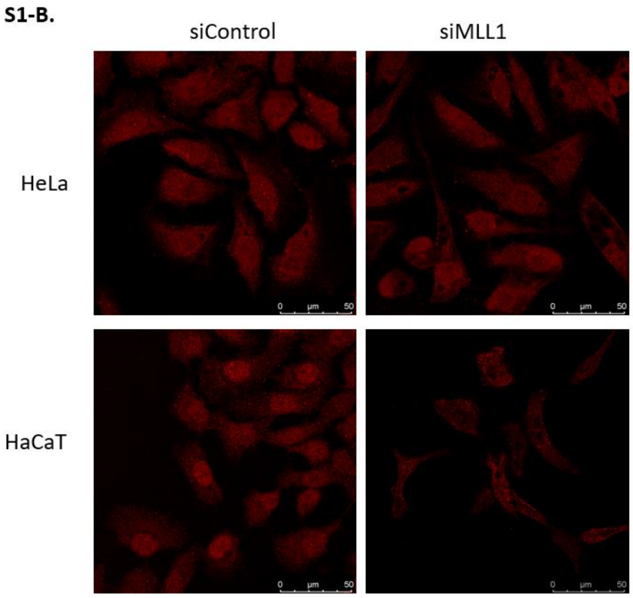


#### Supplementary figure S1:

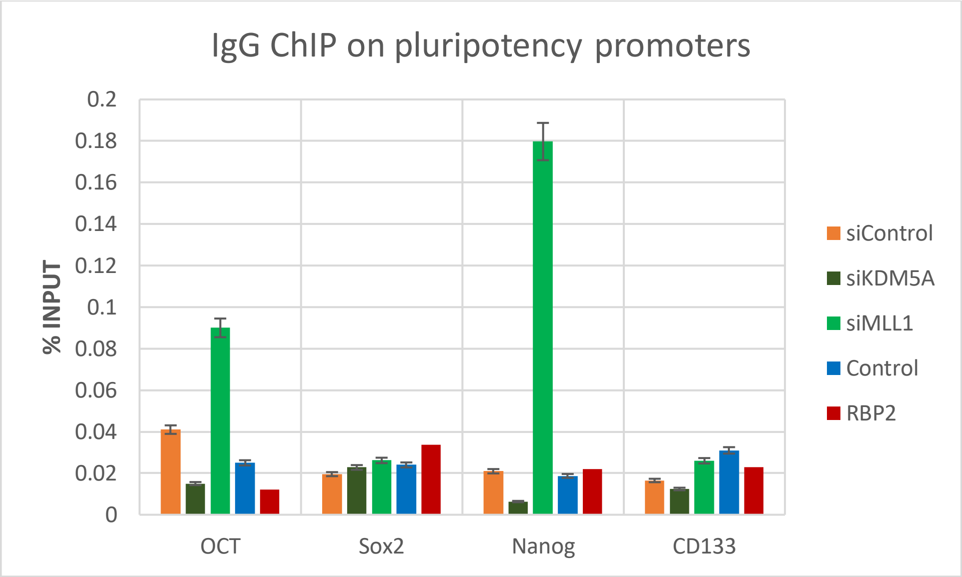


**Supplementary figure S2**

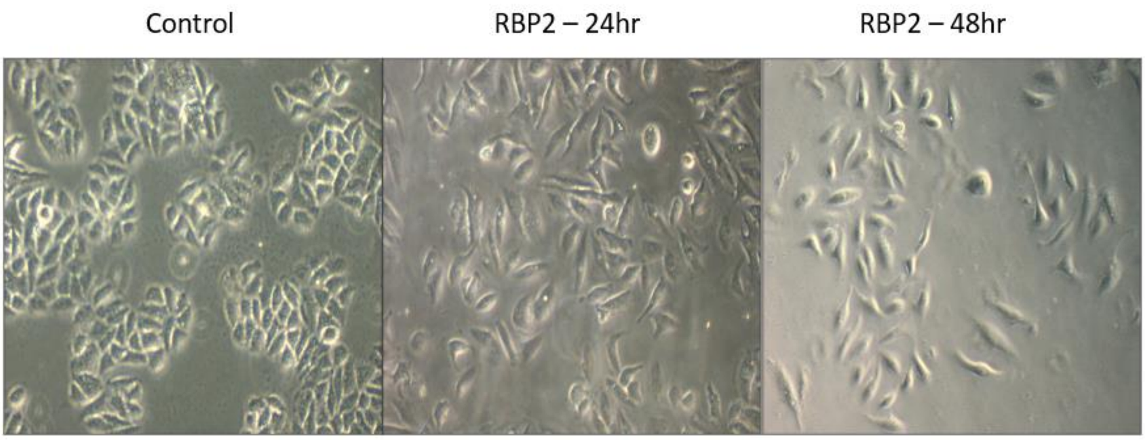


**Supplementary figure S3:**

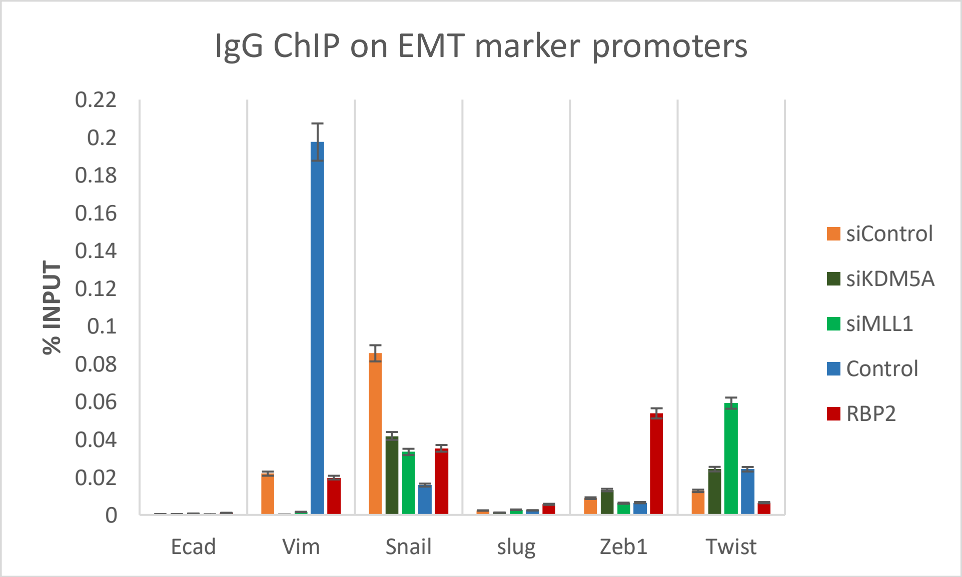

**Supplementary figure S4:**

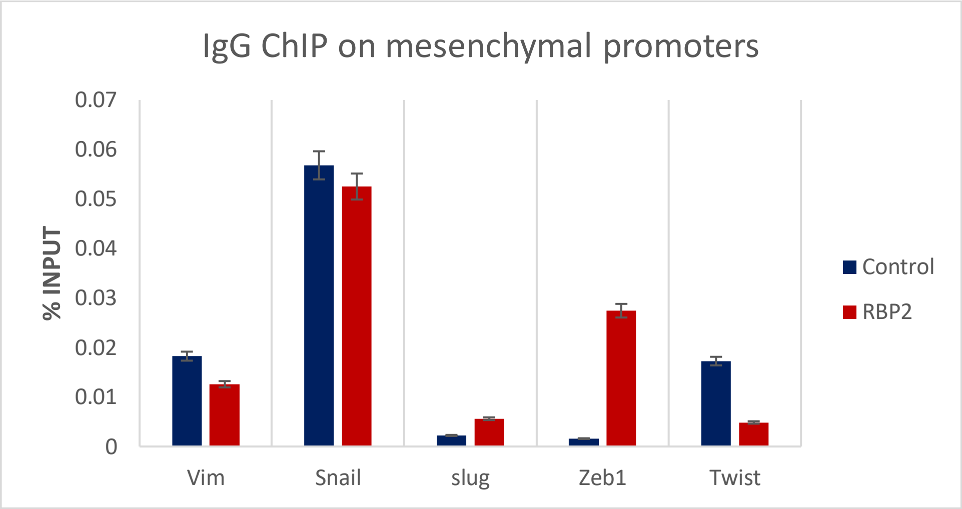


**Supplementary figure S5**

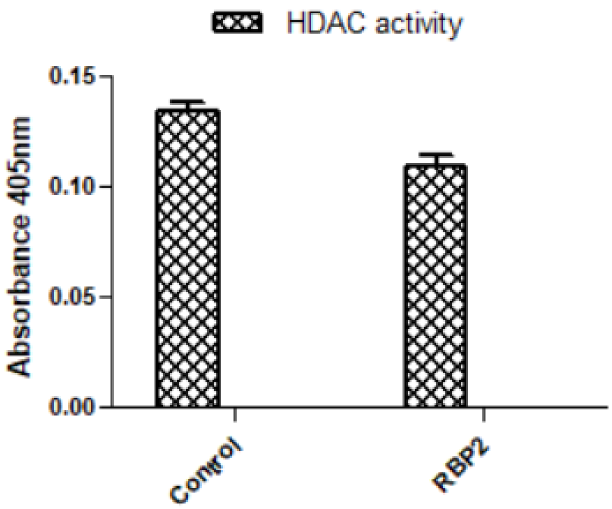


**Supplementary figure S6:**

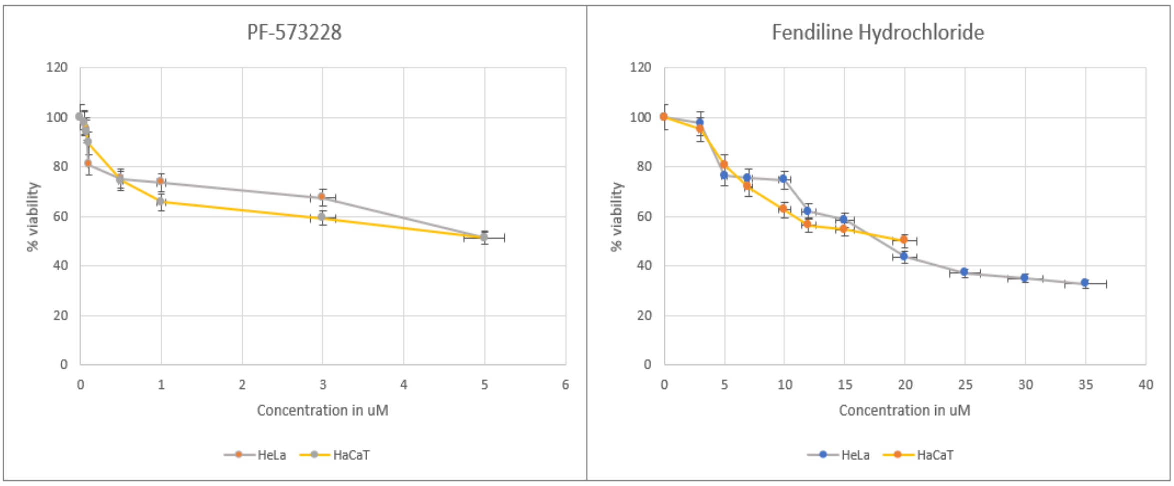


**Supplementary figure S7:**

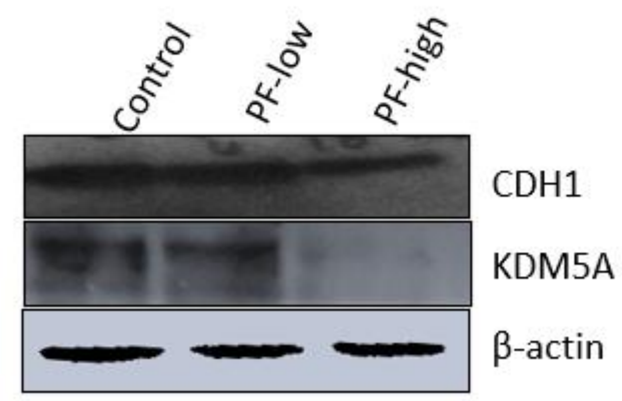


**Supplementary figure S8:**

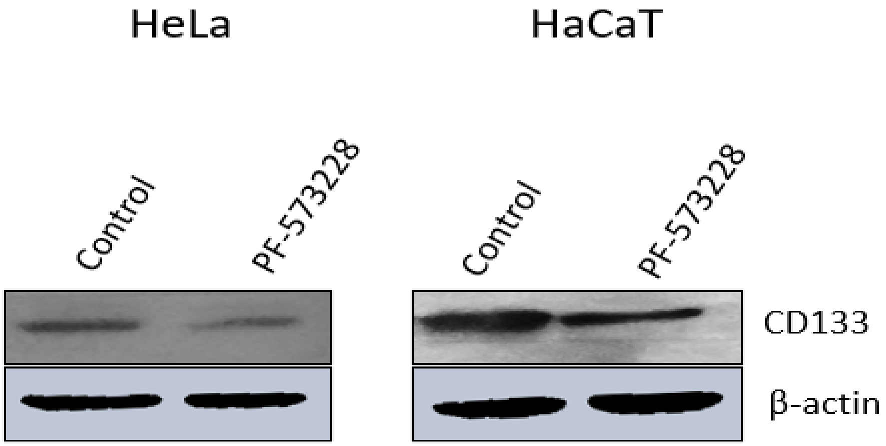


**Supplementary figure S9:**

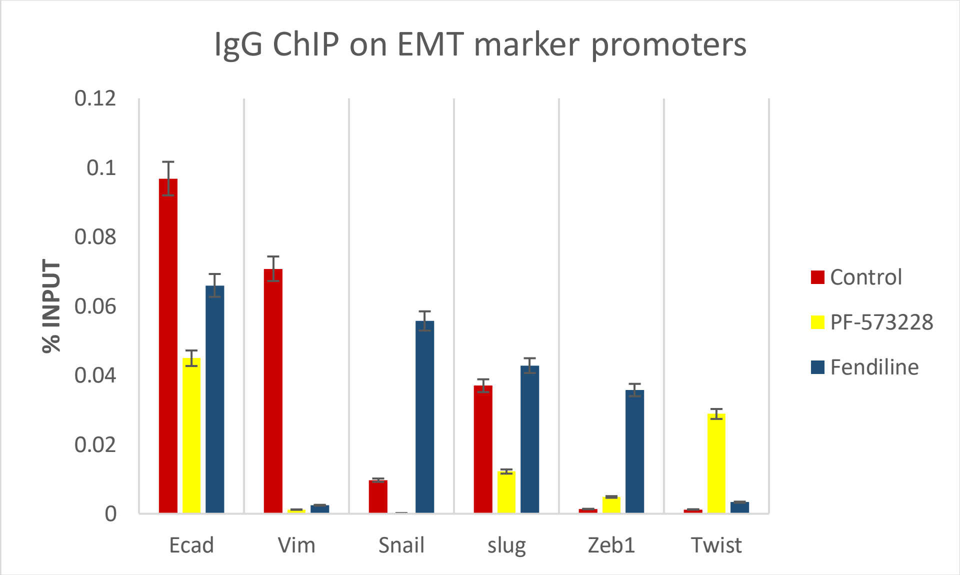


**Supplementary figure S10:**

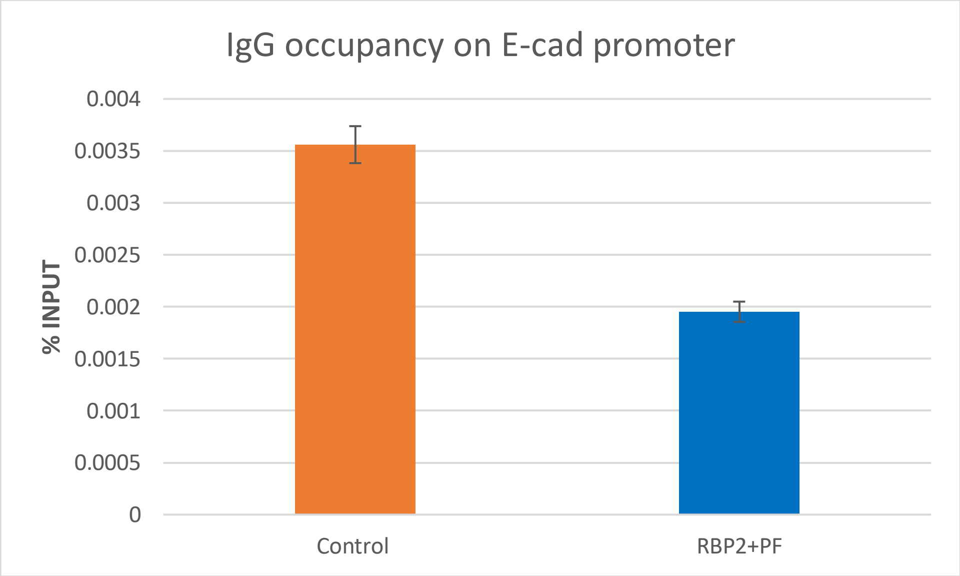


**Supplementary figure S11:**

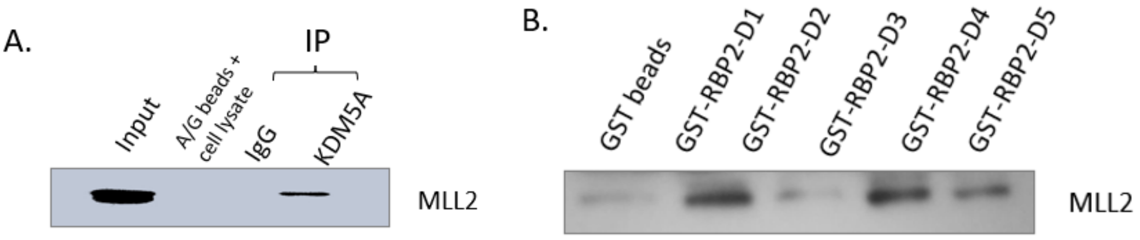


**Supplementary figure S12:**

→ Western Blots : Raw data.

Fig 2A.1 (Hela)

Ecad

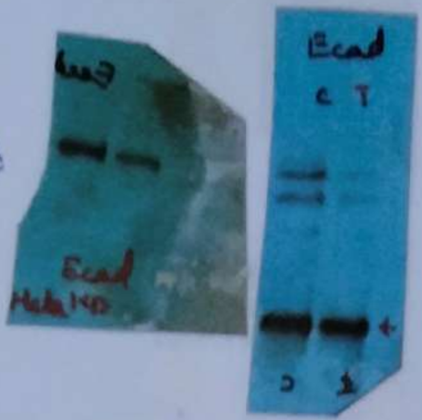


CD133

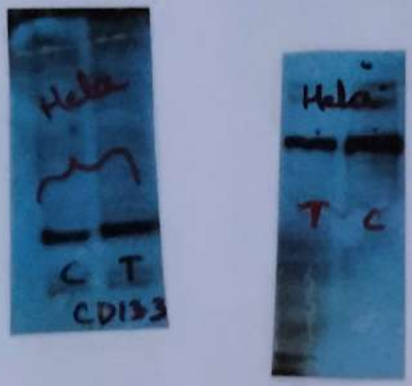


OCT4

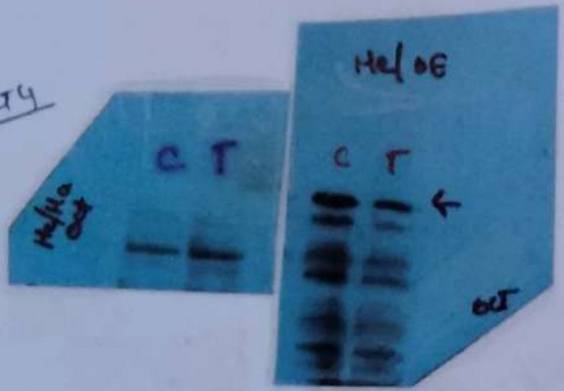


SOX2

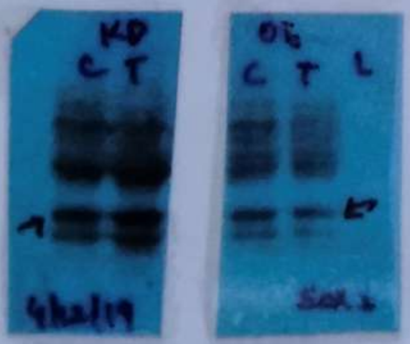


KLF4

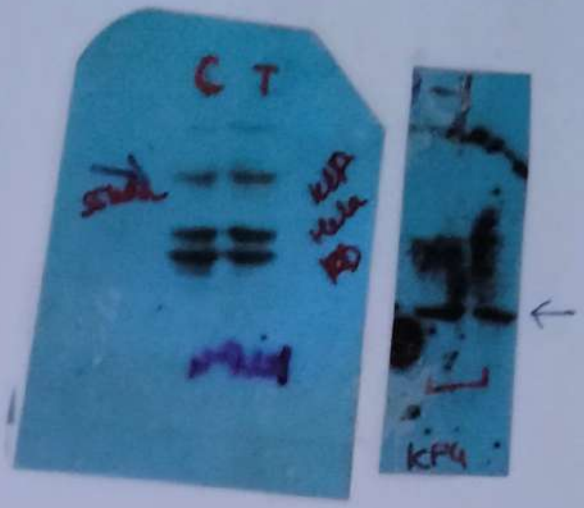


BA

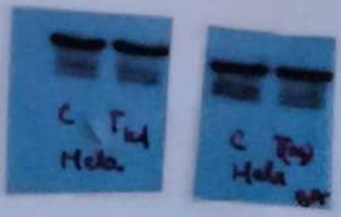

Fig 2A.2 (HaCaT)

Ecad

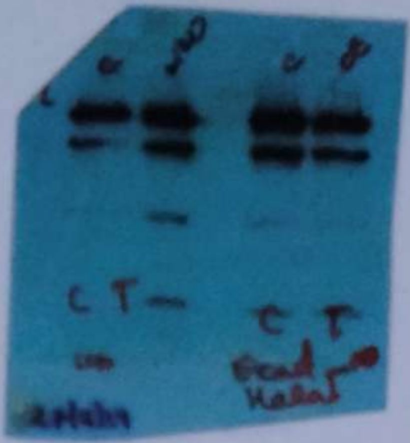


CD133

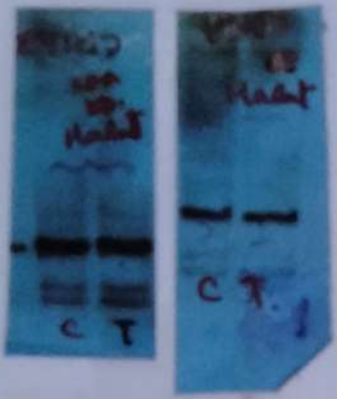


OCT 4

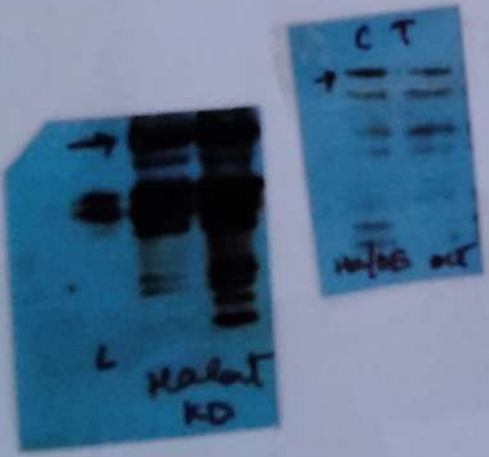


SOX2

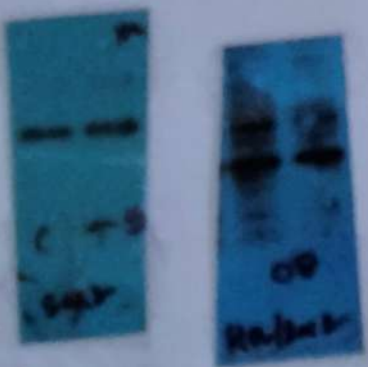


KLF4

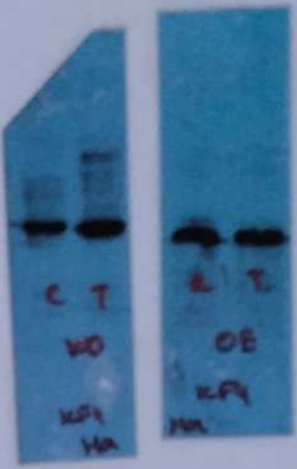


BA

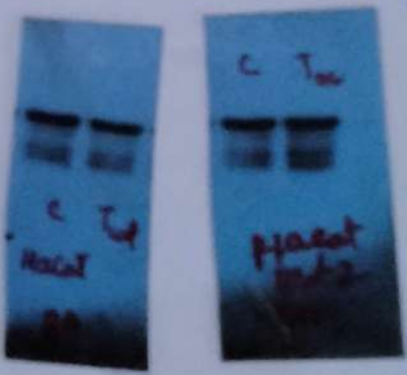

Fig 2A.3 (PC3)

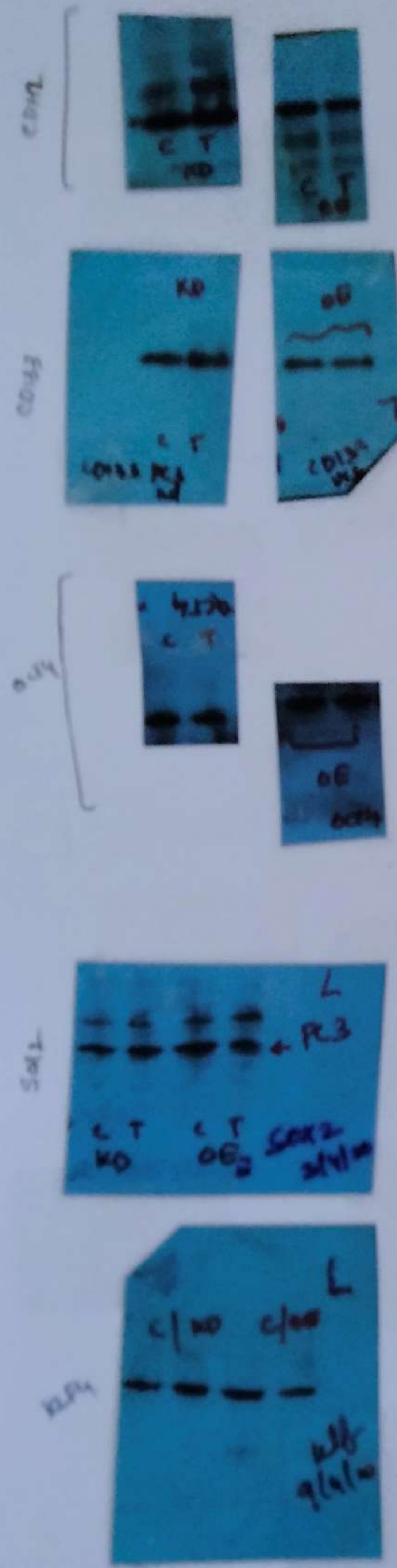


BA.

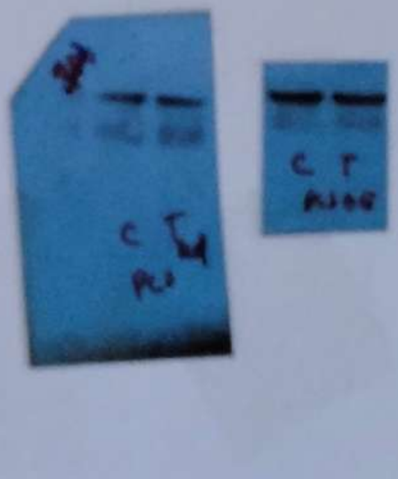

Fig 5.A (Hela)

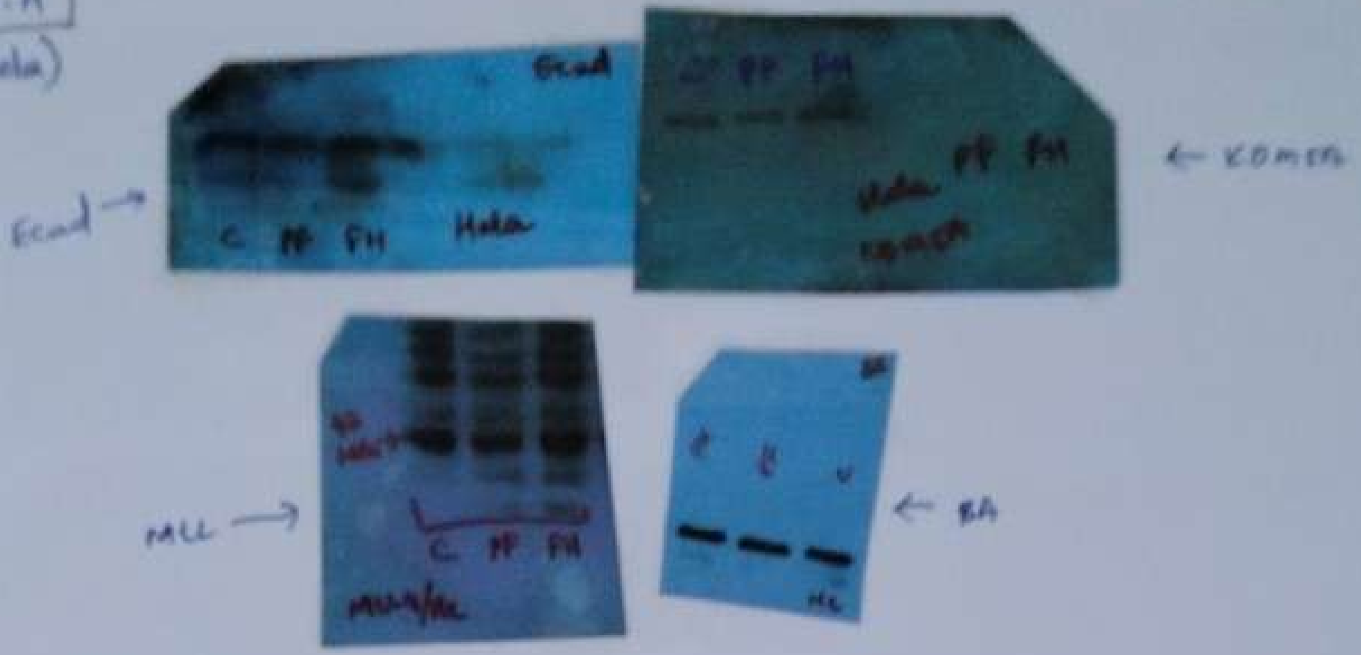


Fig 5B (HaCaT)

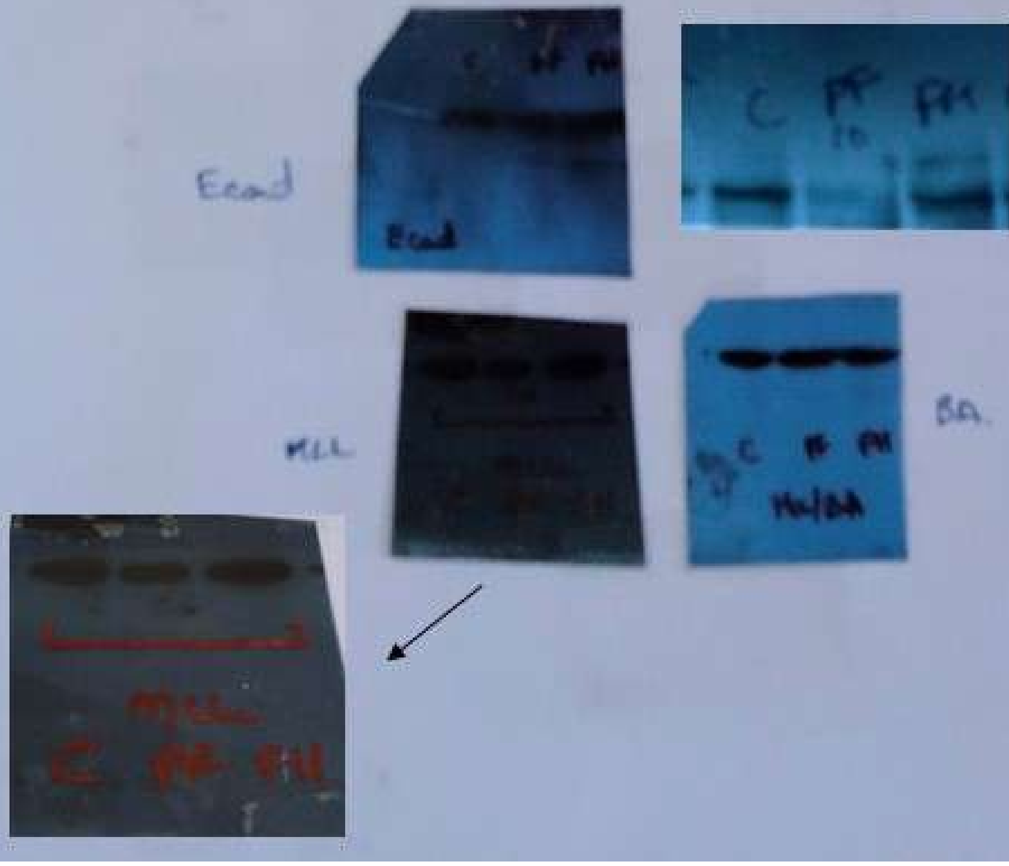

Fig 7A

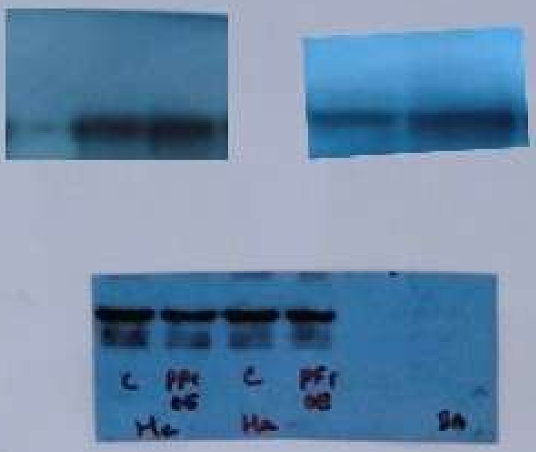


Comp

BA

Fig 8. Co-IP & Pulldown in U87MG cell line :

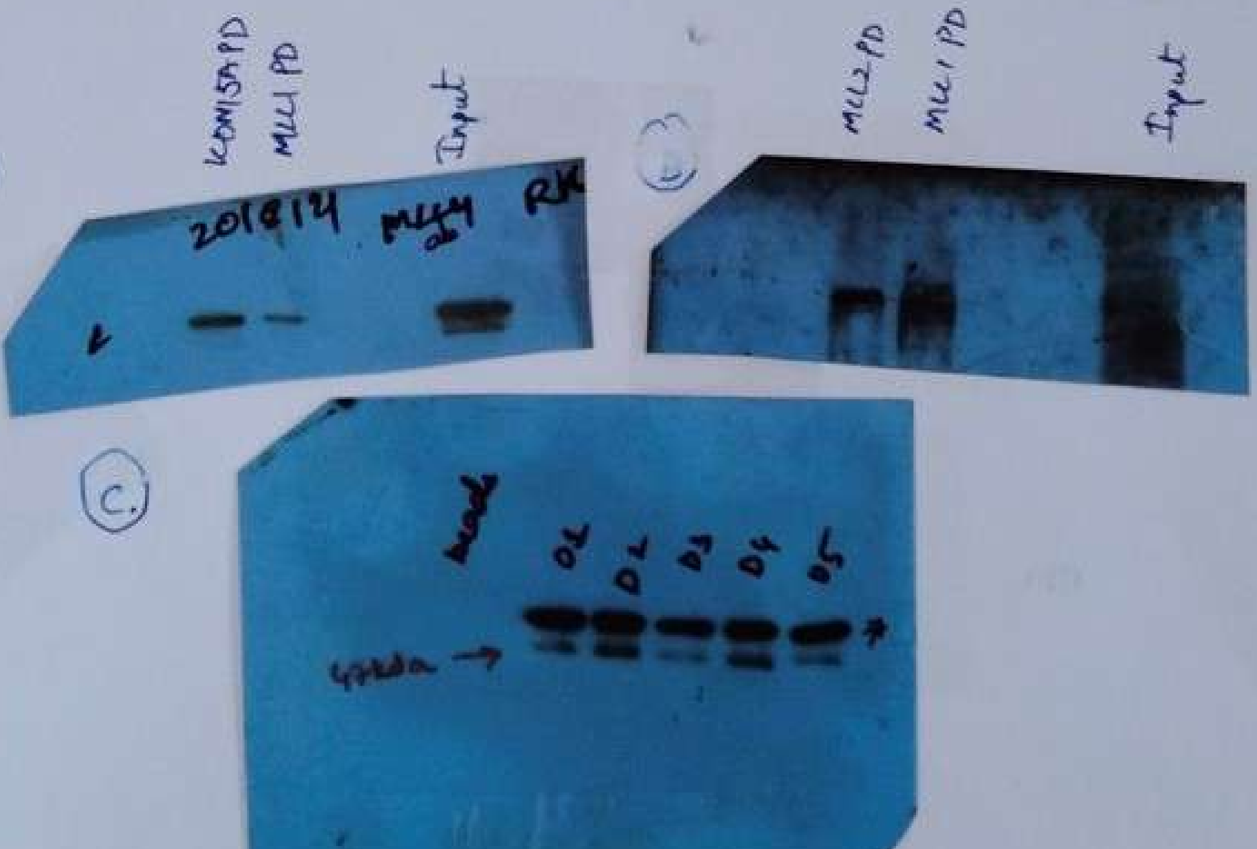


* Non-Specific bands were detected during Pulldown experiments.

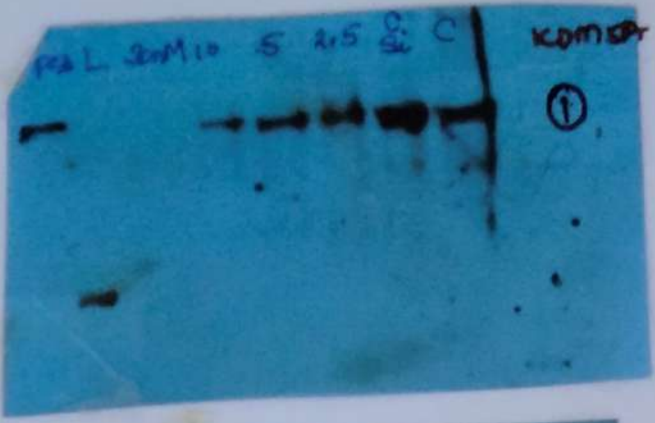


Supp fig 1 . A.
Hela KDM5A - KD : KDM5ab.

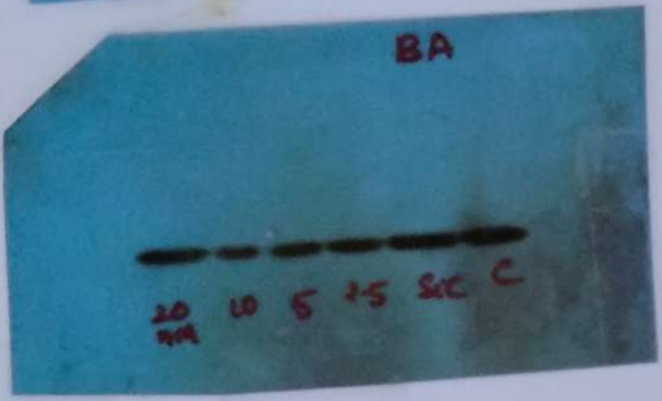


1.A
Hela KDM5A-KD : B- actin ab

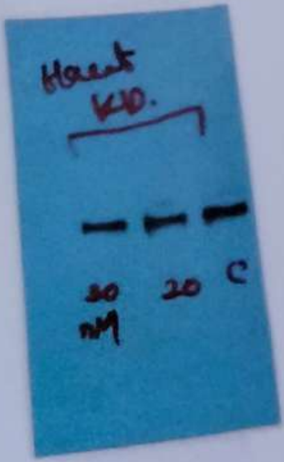


1B.
HaCaT KDM5A-KD : - KDM5 ab

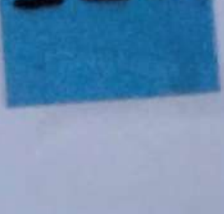

1B.
HaCaT KDM5A - KD : B- actin ab.

1C.

PC3 - KDM5A-KD :
KDM5 ab.

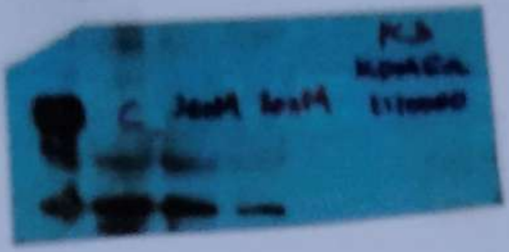

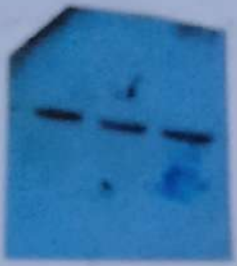

SP-(2)

Supp fig 1D.

Hela KDM5A - OE - KDM5ab.

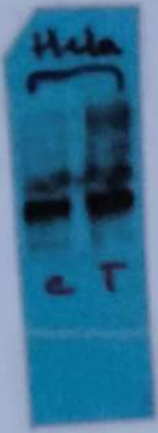


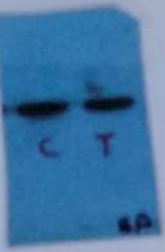


1E.

HaCaT KDM5A - OE - KDM5ab.

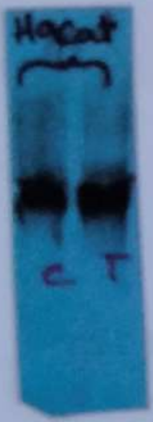


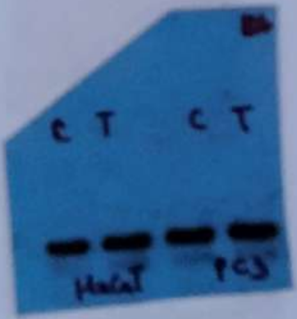


1F.

PC3. KDM5A - OE - KDM5ab

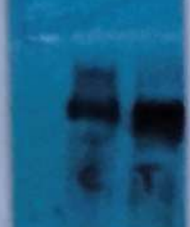

Supp fig. 8 (pg 27)

### PF-573228 - low & high treatment.

Blotted with E-cad ab

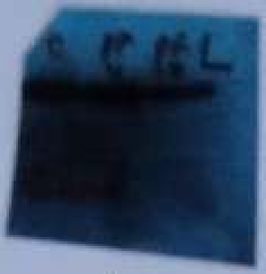


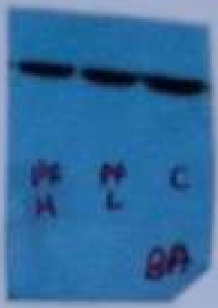


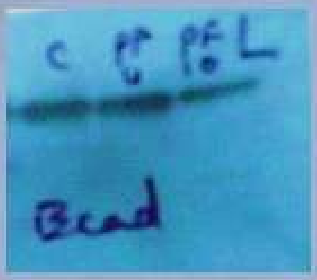


Please note that this blot isn't contrast adjusted, but as X-ray developed was very dark (the blot above) and couldn't be normally scanned, it was scanned against bright background.

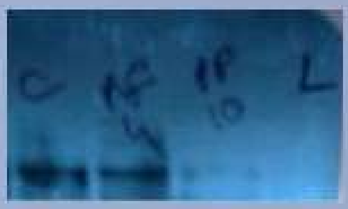


KDM5A

SP - 3

Supp fig 12.

Co-IP & Pulldown in HaCaT cell line:

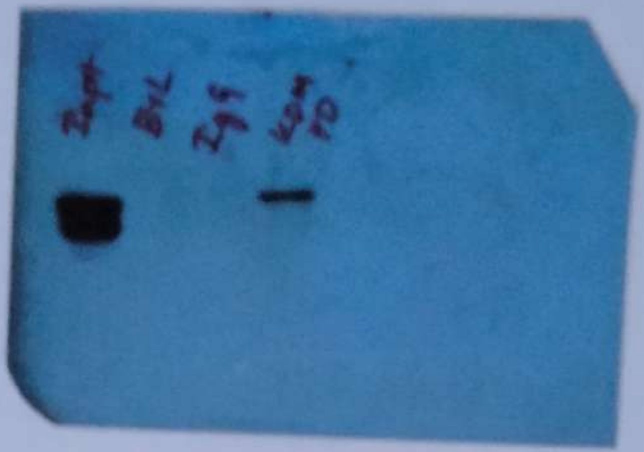

A.

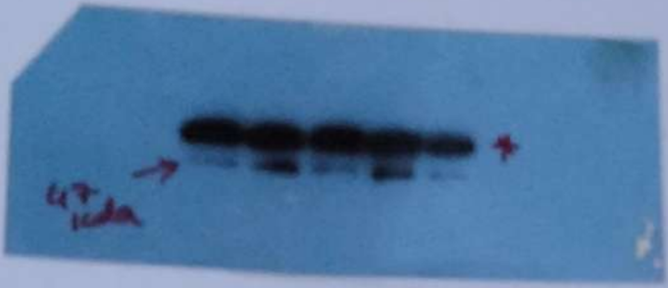


B.

* Non-Specific bands during PD-expt.
